# Haplotype Bias Detection Using Pedigree-Based Transmission Simulation: Traces of Selection That Occurred in Apple Breeding

**DOI:** 10.1101/2025.03.16.643583

**Authors:** Hideto Mochizuki, Mai F. Minamikawa, Kosuke Hamazaki, Miyuki Kunihisa, Shigeki Moriya, Koji Noshita, Takeshi Hayashi, Yuichi Katayose, Toshiya Yamamoto, Hiroyoshi Iwata

**Affiliations:** Laboratory of Biometry and Bioinformatics, Department of Agricultural and Environmental Biology, Graduate School of Agricultural and Life Sciences, The University of Tokyo, 1-1-1 Yayoi, Bunkyo, Tokyo 113-8657, Japan; Institute for Advanced Academic Research (IAAR), Chiba University, 1-33 Yayoi, Inage, Chiba, Chiba 263-8522, Japan; Molecular Informatics Team, RIKEN Center for Advanced Intelligence Project (AIP), RIKEN, 178-4-4 Wakashiba, Kashiwa, Chiba 277-0871, Japan; Institute of Fruit Tree and Tea Science, National Agriculture and Food Research Organization (NARO), 2-1 Fujimoto, Tsukuba, Ibaraki 305-8605, Japan; Institute of Fruit Tree and Tea Science, NARO, 92-24 Nabeyashiki, Shimokuriyagawa, Morioka, Iwate 020-0123, Japan; Department of Biology, Kyushu University, 744 Motooka, Nishi-ku, Fukuoka, 819-0395, Japan; Research Center for Agricultural Information Technology, National Agriculture and Food Research Organization (NARO), 1-31-1 Kannondai, Tsukuba, Ibaraki 305-0856, Japan; Institute of Crop Science, National Agriculture and Food Research Organization (NARO), 2-1-2 Kannondai, Tsukuba, Ibaraki 305-8518, Japan

**Author notes:** The official email addresses of other authors: Hideto Mochizuki^1^, Mai F. Minamikawa^2^, Kosuke Hamazaki^3^, Miyuki Kunihisa^4^, Shigeki Moriya^5^, Koji Noshita^6^, Takeshi Hayashi^7^, Yuichi Katayose^8^, Toshiya Yamamoto^4^.

## Abstract

With the increasing ability to integrate pedigree and genomic data, it is essential to evaluate their potential to uncover valuable genetic insights that can drive the advancement of crop breeding and conservation of genetic diversity. Pedigree analysis remains a fundamental approach for investigating the inheritance of phenotypic traits, exploring evolutionary history, and understanding hybridization processes in crop plants. Among these approaches, gene drop simulations using pedigree and allele origin data enable the construction of genetic maps and provide insights into complex genetic backgrounds. In this study, we developed a new method to identify useful genetic regions associated with single-nucleotide polymorphism (SNP) markers based on gene drop simulations, focusing on 185 Japanese domestic apple cultivars. By performing 10 million gene drop simulations, we generated null distributions for each founder haplotype, which revealed SNP markers with significant frequency biases, which is a potential signal for selection. Frequency biases were identified in eight founder haplotypes that were particularly consistent with genome-wide association studies peaks associated with key fruit traits such as malic acid and fructose content. Gene Ontology enrichment analysis suggested that these SNPs are not only associated with fruit traits but may also play a role in critical biological functions, including stress tolerance and reproductive processes, highlighting their broader relevance to crop resilience. Our integrative approach, which combines founder haplotype analysis with extensive gene drop simulations, effectively detects selection pressure, provides new insights into the genetic basis of apple breeding, and identifies SNP markers with strong potential to improve breeding programs.

## Introduction

Historical breeding materials are of paramount importance for perennial plants such as fruit trees, as they have accumulated rich genetic variation over many years^1^. Woody fruit trees, such as apples, are often self-incompatible, leading to well-documented pedigree information compared with crops, such as rice, which have more complex genetic backgrounds owing to selfing^2, 3^. Therefore, data-driven breeding approaches that use pedigree information allow for the more effective identification and use of beneficial traits^4^.

In recent years, advancements in sequencing technology have made it possible to sequence genomes quickly and affordably, extending their capabilities to plant breeding^5^. For example, genome-wide association studies (GWAS) ^6^, which identify candidate genes responsible for traits, such as disease resistance, have been applied to various plant species, including apple^7, 8, 9^, rice^10^, wheat^11^, maize^12^, and citrus^13^. This demonstrates that GWAS have become a mainstream method in modern plant breeding. Furthermore, in breeding materials with complete pedigree and single-nucleotide polymorphism (SNP) information, founder haplotypes can be visualized and automatically traced^7^, enabling breeders to use historical data more effectively.

In a study conducted by Minamikawa et al.^7^, gene drop simulations^14^ were used to validate the non-random transmission of founder haplotypes detected in GWAS. However, the bias detected using this method is not limited to markers significantly associated with the GWAS peaks. In this study, we divided the Japanese domestic apple population based on pedigree information visualized using Helium^15^ and expanded gene drop simulations on generational and genome-wide scales. This approach allowed the identification of potential biases in the frequencies of founder haplotypes by comparing them with random transmissions across all markers. By examining genome-wide changes in founder haplotype frequencies without focusing on specific traits, we inferred the occurrence of intended and unintended selections based on these changes. Additionally, the temporal dynamics of these founder haplotypes were tracked to provide insights into breeding trends during specific periods.

To the best of our knowledge, no previous studies have conducted whole-genome gene drop simulations on apples or detected biases in founder haplotypes across markers in other plant species using this method. This study builds on the work of Minamikawa et al. ^7^ by investigating the underlying reasons for the selection of biased markers and founder haplotypes across the entire genome. Specifically, we compared biased markers with those identified by the GWAS. In cases where there was an overlap between biased markers and GWAS peaks, we analyzed generational changes in the trait, the effect of each founder haplotype on the trait, and frequency changes of the founder haplotypes at the marker to infer the reasons for selection. When no overlap was observed, it suggested the possibility of selection for traits other than those related to fruit characteristics. To explore this further, we combined gene drop simulations with gene ontology (GO) enrichment analysis^16^ to gain insight into the reasons for selection beyond the traits targeted by the GWAS. This study employed a quantitative approach to evaluate the impact of breeding-induced biases on gene function and assessed genes by integrating GWAS and GO enrichment analyses.

Previous studies have developed gene drop simulations^14^ using pedigree and allele information, which have been used to construct genetic maps and understand genetic backgrounds^17, 18^. In apples, a similar gene drop simulation was performed using founder haplotypes^19^ instead of allele information^20^ to validate the non-random transmission of founder haplotypes detected in GWAS^7^. However, the biases detected using this method are not limited to markers that are significantly associated with GWAS peaks. It is also important to consider when these biases arise because they may reflect the preferences of consumers and breeders at particular times. To address this, we extended the gene drop simulation to the whole genome and ran it across all generations based on pedigree data visualized using Helium^15^. This allowed us to detect biases independent of the GWAS results. By integrating this approach with GO enrichment analysis^16^, we gained insights into selective pressures that may extend beyond the traits traditionally targeted by GWAS.

## Results

### Detecting frequency bias of founder haplotypes using gene drop simulation

To generate a null distribution for the 14 founder haplotypes derived from the seven apple founder cultivars, 10 million gene drop simulations were performed (Supplementary Fig. 1). Two types of null distribution were created: one for the entire population and one for each generation. This generation constitutes a division of the genealogical information inscribed using Helium and approximates the age at which the variety was created. SNP markers with significantly altered frequencies for each founder haplotype were detected by comparing the observed founder haplotypes with null distributions. Frequency bias was identified in eight founder haplotypes (1, 2, 3, 4, 5, 6, 8, and 13) when the entire population was tested (Fig. 1; Supplementary Table.). In tests for individual generations, significant changes in the frequencies of some founder haplotypes were observed starting from the third generation (Supplementary Figs. 2–6). SNP markers detected in the separate-generation tests were identical to those found in the entire population. Hereafter, the results focus on the population-wide test, which is considered to have higher statistical power.

**Fig. 1.**
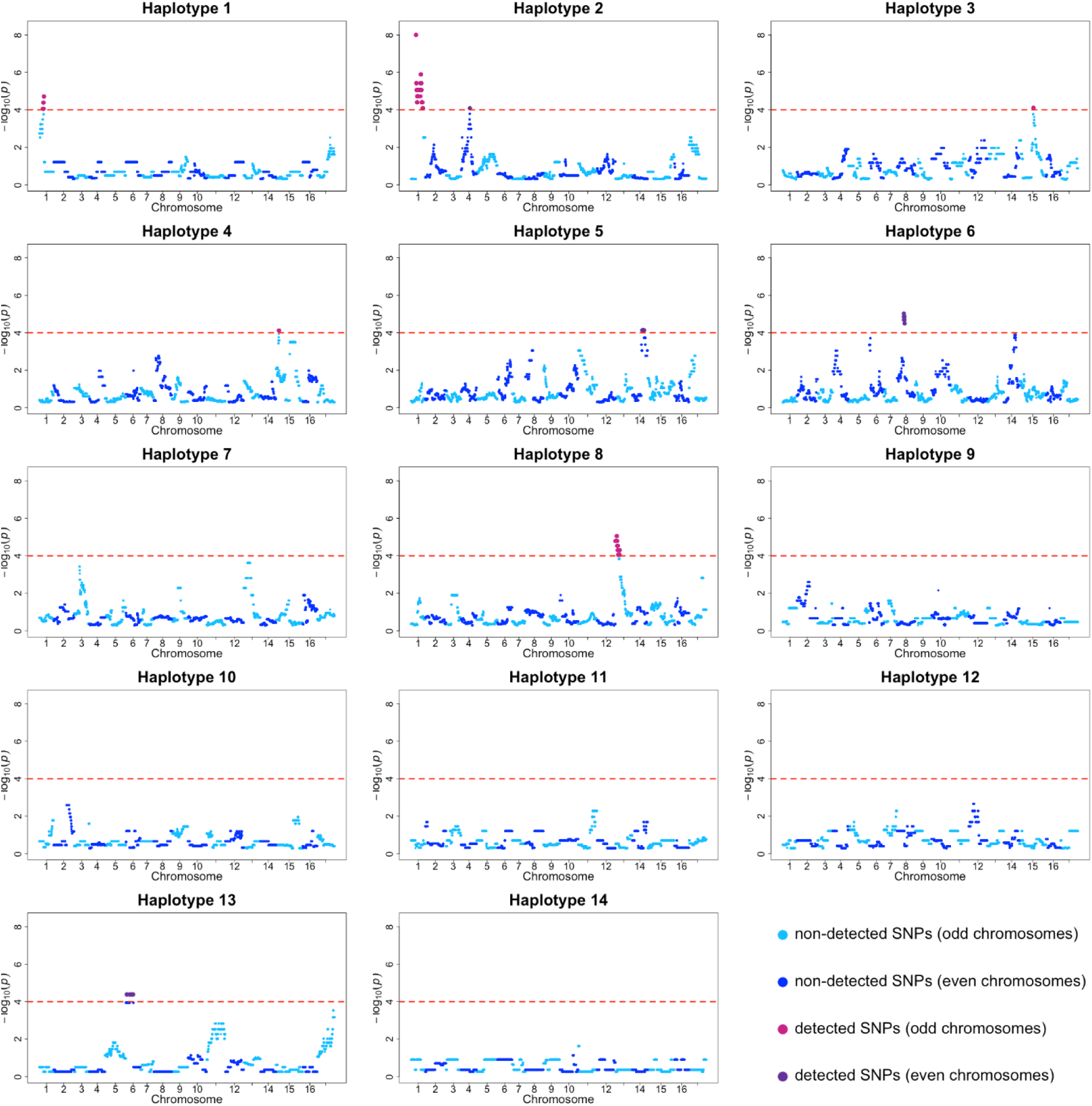
Detection of frequency bias in the 14 founder haplotypes using gene drop simulations. Regions considered to be significantly selected for each founder haplotype are shown. The horizontal axis represe ts the genomic position of SNPs, and the vertical axis shows the value from the simulations. Purple dots indicate significant regions, while blue dots represent non-significant regions. The red dashed lines mark the threshold ().

### Concordance between SNP markers with biased frequencies of founder haplotypes and previously reported GWAS peaks or reference annotation data

To investigate the reasons behind the changes in founder haplotype frequencies, we compared SNP markers located on the coding DNA sequence that significantly altered founder haplotype frequencies with previously reported GWAS peaks for fruit traits. The SNP markers with significantly altered founder haplotype frequencies coincided with the GWAS peaks for 11 fruit traits (Supplementary Table). SNP markers were grouped according to their corresponding GWAS target traits and loci (Supplementary Table). Subsequently, the markers were divided into 12 blocks based on chromosome and physical position, and the SNP marker with the highest -log_10_(*p*) value was designated as the representative marker for each block (Supplementary Table; red text cells in the block column). The founder haplotypes that were significant for these markers were then plotted to show their frequency changes over generations, the estimated effect of each founder haplotype on the marker, and the generational evolution of fruit traits associated with the matched GWAS peaks.

The results showed that among the 12 blocks, seven exhibited changes in the frequency of the founder haplotype of interest, with consistency between this change and the phenotypic value transition (Supplementary Figs. S7, S9–13, and S16). In contrast, three blocks did not show this consistency (Supplementary Figs. S8, S14, and S15). The two blocks displayed no changes in the relevant phenotypic trends across generations (Supplementary Figs. S17 and S18). For example, the phenotypic value of malic acid tended to decrease over time, but the SNPs associated with it increased significantly in frequency, showing a negative effect in all cases (Supplementary Figs. S10–12).

In addition to the above analysis, we compared the annotations assigned to the ‘Golden Delicious’ (GDDH13) reference to explore factors that may have contributed to the selection of these regions beyond fruit traits. This comparison revealed that 68 SNP markers matched 35 genes within the SNP markers identified as founder haplotype biased that associated with previously reported GWAS peaks (Supplementary Table.). These SNP markers may play roles in processes beyond fruit traits, such as cell division, pollen development and germination, drought and salt stress tolerance, reproduction, and the maintenance of vital activities.

Moreover, the annotation of apple genes in the gene regions containing the corresponding SNP markers was examined to determine the reason for the bias in the founder haplotype frequencies of SNP markers that did not match previously reported GWAS peaks. In total, 669 SNP markers with biased founder haplotype frequencies corresponded to 351 gene regions (Supplementary Table). These SNP markers, along with those that matched the reported GWAS peaks, may be involved in plant growth processes, such as growth regulation and RNA binding.

### GO enrichment analysis to estimate the reasons for selected regions

GO enrichment analysis suggested that GO terms in the biological process category were not significantly (*p* < 0.05) enriched within genes containing 669 SNP markers with significantly altered founder haplotype frequencies and unmatched GWAS peaks. GO terms of the Molecular Function (MF) category, ‘transferase activity, transferring phosphorus-containing groups’ and ‘sequence-specific DNA binding,’ were significantly (*p* < 0.05) enriched within the genes (Fig. 2(a)). ‘Sequence-specific DNA binding’ had the lowest *p*-value in the MF category. Additionally, when we analyzed the genes associated with each function in Figure 2(a), *HSF4*, *LOC103416823*, and *WRKY12* genes were each linked to one function (Fig. 2(b)). Within the network of these genes, the RNA polymerase II transcription regulatory region sequence-specific DNA binding appeared to have the greatest downstream function (Supplementary Fig. S19). Furthermore, the cellular component (CC) category was suggested to be related to organelle and nuclear lumens (Supplementary Fig. S20).

**Fig. 2.**
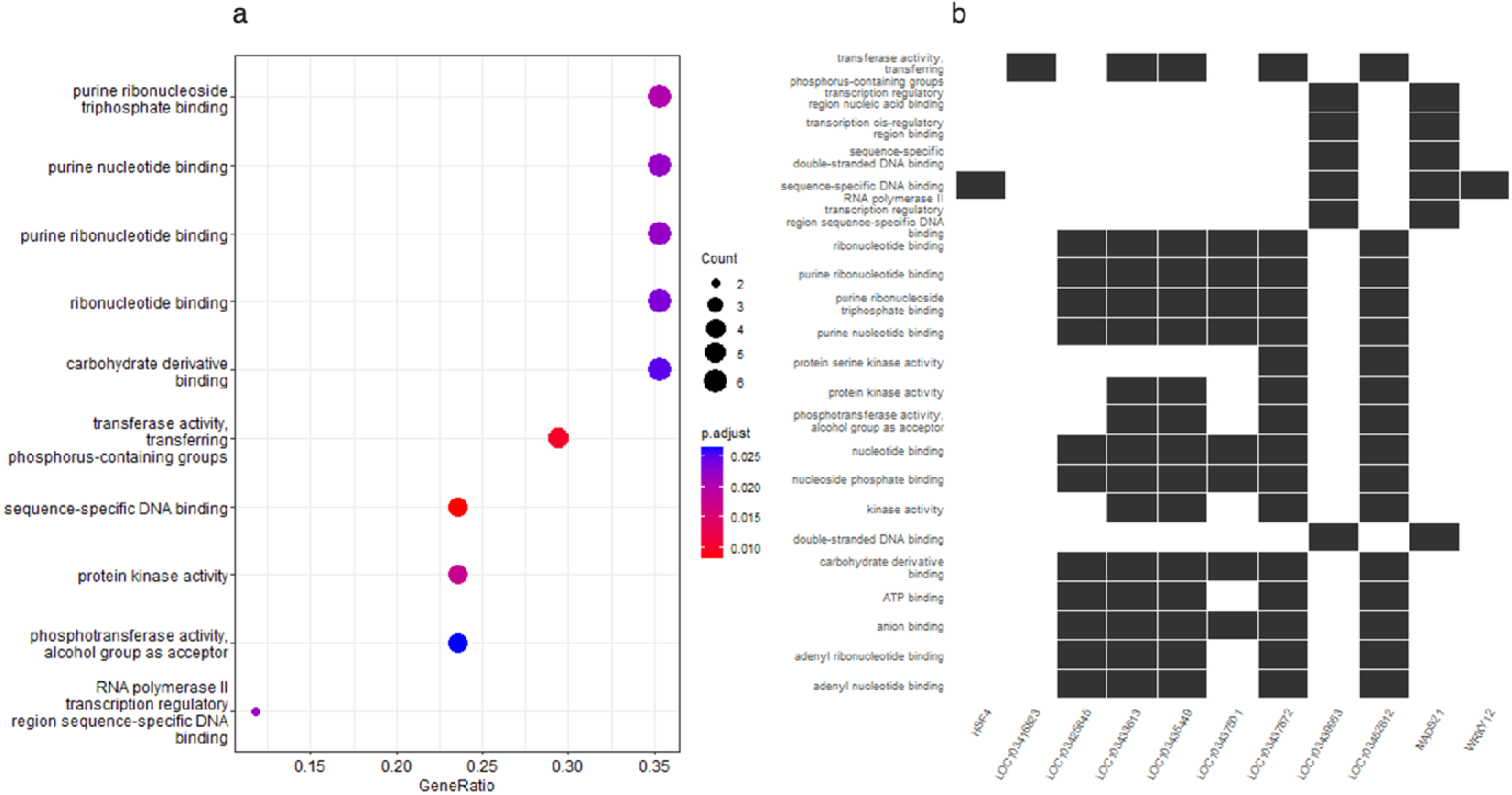
Results of gene ontology enrichment analysis in molecular functions (MF) (a) The gene ontology (GO) term within the MF category enriched in genes containing SNP markers with signific ntly altered founder haplotype frequencies. The horizontal axis represents the proportion of genes associated with each GO term, and the size of each dot represents the number of genes. The color of the dots indicates the *p*-value, with red representing statistically significant associations, implying a stronger likelihood of being linked to that function. (b) A figure showing the genes associated with each GO term. Sequence-specific DNA binding (fifth row from the top), which had the lowest *p*-value, was linked to four genes: *HSF4*, *LOC103439953*, *MADS21*, and *WRKY12*.

## Discussion

In this study, we introduced an approach based on pedigree and gene drop simulations to identify biases in the frequency of founder haplotypes. Since the frequency of founder haplotypes contributing to valuable phenotypes is expected to increase within breeding populations, the regions identified using this method offer valuable insights for breeding. Focusing solely on founder haplotype frequencies might lead to the conclusion that all founder haplotypes of ‘Fuji’ are beneficial, as haplotypes from ‘Fuji,’ a parent with significant contributions to later generations, would naturally have higher frequencies in the breeding population. However, by comparing the degree of bias in founder haplotypes with random inheritance, our new method enables the evaluation of their contributions to later generations. Unlike traditional methods, such as GWAS and genomic prediction, which rely heavily on phenotypic data, our method does not depend on such data, making it robust and versatile.

Previous research has often traced founder haplotypes at a single locus^7^ or focused on genetic diversity using gene drop simulations^14^; however, a genome-wide method for identifying useful genomic regions through gene drop simulations is lacking. This study aimed to investigate the causes of changes in founder haplotype frequencies at certain SNP markers by tracing them across the whole genome, generation by generation. Additionally, we analyzed the gene regions to determine the possibility that selection may have occurred for unmeasured traits. Therefore, the proposed method offers a unique approach.

By investigating the functions of the SNP markers with significantly altered founder haplotype frequencies detected using our method, we aimed to determine the reasons for selecting these regions. For example, in Supplementary Fig. S7 (a) and (b), the founder haplotype 6 from ‘Golden Delicious’ showed a significant change in frequency at the SNP marker SNP_FB_1117728, which coincided with the GWAS peak for fructose. The founder haplotype 6 in this region exhibited a strong positive effect on fructose content, with its frequency increasing significantly above the simulation threshold by the fourth generation (Supplementary Fig. S7 (c)). This suggests that the founder haplotype 6 at SNP_FB_1117728 from ‘Golden Delicious’ may have been selected to increase fructose content. Given that fructose is sweeter than other sugars^21^ and consumer preference often leans toward sweeter varieties^22^, it is possible that consumer choice may have influenced the selection for increased fructose content.

Furthermore, GO enrichment analysis using genes containing SNP markers with significantly altered founder haplotype frequencies, GO terms of the MF category, such as ‘transferase activity, transferring phosphorus-containing groups’ and ‘sequence-specific DNA binding,’ were identified (Fig. 2). Specifically, focusing on sequence-specific DNA binding, four genes, *HSF4*, *LOC103439953*, *MADS21* and *WRKY12* -- were found to be involved in regions with significantly altered founder haplotype frequencies. *HSF* plays an important role in flavonoid biosynthesis and drought resistance^23^. Additionally, *HSF4* is regulated by cold stress in bananas^24^, suggesting its potential role in cold and heat tolerance in apples.

*MADS21* regulates unsaturated fatty acids in the palm by contributing to metabolic genes^25^ and is involved in the regulation of fat development and flowering transition in Arabidopsis^26, 27, 28^,. The *WRKY* family is known for its role in abiotic stress responsiveness, including resistance to *Alternaria alternata* in apple^29^. Additionally, *WRKY12* exhibited the opposite regulatory effect on flowering under short-day conditions^30^. This implies that the regions detected using this method may have undergone significant changes in founder haplotype frequencies owing to selection for traits such as growth, disease resistance, and environmental adaptation, which are often not measured in phenotypic assessments.

Unconscious selection by breeders, reflected in regions with significantly altered founder haplotype frequencies, likely led to an increase in the number of haplotypes associated with beneficial effects on physiological functions. Unconscious selection in breeding processes has rarely been explored, making our methodology a pioneering contribution to our understanding of the historical context and unintentional decisions made during the breeding of apple varieties.

In recent years, serious diseases such as *Alternaria* fungus^31^ and burn diseases have had a significant impact on apple breeding. For example, among the seven ancestral Japanese apple varieties (Founder cultivar; ‘Ralls Janet’, ‘Delicious’, ‘Golden Delicious’, ‘Jonathan’, ‘Worcester Pearmain’, ‘Indo’, ‘Cox’s Orange Pippin’)^17^, ‘Indo’ and ‘Delicious’ are particularly susceptible to *Alternaria alternata*, and this low resistance has been inherited by many of their offspring. Additionally, ‘Fuji’ is prone to burn, a disease that often causes wilting^32^. Marker-assisted selection is available for these diseases^33^. However, although such methods are highly accurate, they are labor-intensive for the comprehensive identification of resistance genes. Our method may not only allow for the detection of gene regions potentially associated with resistance to such diseases but also provide a comprehensive approach to identifying useful gene regions that might be useful for resistance against severe diseases that have not yet been extensively studied, potentially opening new possibilities for future breeding efforts. Furthermore, this method is not limited to apples and is applicable when the pedigree is limited and founder haplotype information is complete. This adaptability makes it a valuable tool for other fruit tree species such as citrus^34^ and peach^35^.

In conclusion, our method, based on genealogical information and gene drop simulations of founder haplotypes, is a valuable tool for detecting biases in founder haplotype frequencies, while minimizing the influence of breeding parents with substantial contributions to subsequent generations, such as ‘Fuji.’ By comparing these detected biases with the GWAS peaks, we inferred the selective intentions behind the selection of specific regions, offering insights into the often-unexplored realm of breeder decisions. Additionally, by comparing regions with significantly altered founder haplotype frequencies with annotated regions, we identified useful genetic regions that may not have been directly targeted by breeding programs. Because our method does not require phenotypic measurements, it allows the exploration of a wider range of valuable genetic regions, providing new opportunities for future breeding efforts.

## Materials and Methods

### Plant materials

Data obtained from 185 domestic Japanese apples (*Malus domestica* Borkh.) varieties^7^ were used in this study. These varieties originated from seven founder cultivars: ‘Ralls Janet’, ‘Delicious’, ‘Golden Delicious’, ‘Jonathan’, ‘Worcester Pearmain’, ‘Indo’, ‘Cox’s Orange Pippin’. All the materials mentioned above were cultivated at the Apple Research Station, Institute of Fruit Tree Science, NARO, Japan.

### Pedigree information, genotypes, and phenotypes

Pedigree information for 185 domestic Japanese apple varieties was used in this study. SNP genotypes for 11,786 markers (Illumina Infinium array) were obtained, and sporadic missing SNP genotypes were imputed using Beagle ver. 4.0^21^, as reported by Minamikawa et al.^7^. The SNP data of the varieties were characterized by 14 founder haplotypes derived from seven founder cultivars, as detailed in the same publication. The phenotypic data for 23 fruit traits evaluated over multiple years^7^ were included in this study.

### Gene drop simulation to detect the frequency bias of founder haplotypes

The genome-wide haplotype composition of a population reflects the effects of selection and mating during breeding, particularly around gene regions related to the target trait. This study focused on the usefulness of pedigree-based gene drop simulation^10^ to account for bias created by breeding parents. We combined this approach with founder haplotype information that represents the origin of SNP alleles. The aim of this study was to identify regions that have undergone intended and unintended selection during the breeding process. Because the apples used in this study were diploid (2n = 2x = 34), this gene drop simulation assumed the null hypothesis that one of the two founder haplotypes at an SNP locus was randomly selected with a probability of 1/2 in subsequent generations. Therefore, the transmission of the founder haplotype at a locus was equivalent to performing a Bernoulli trial with *p*= 1/2. In this study, multiple founder haplotypes were assumed, and the frequency distribution of the founder haplotypes in the later population followed a multinomial distribution instead of a binomial distribution as follows:

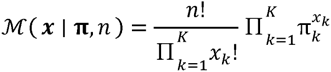

where x= (*x*_1_,*x*_2_,.., *x_k_*) is the number of times about founder haplotype has been obtained, π= (π_1_,π_2_,.., π_k_) is the probability of obtaining a combination of founder haplotypes, *n* is the number of haplotypes (2x population size), and *k* is the number of possible combinations of founder haplotypes in the population. where *π* and x satisfy 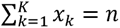, 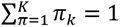. Although it is possible to obtain the frequency of founder haplotypes analytically, as described above, it is difficult to do so for all individuals in this study’s apple population, which consisted of 14 founder haplotypes and 185 parental varieties. To address this, we simulated the spread of founder haplotypes in later generations by creating empirical frequency distributions of founder haplotypes using 10 million gene drop simulations based on pedigree information. This simulation assumed that founder haplotypes were obtained randomly when breeders did not select them as a null hypothesis. This empirical frequency distribution was used as the null distribution. The *p*-value of the observed founder haplotype frequency in each genetic region was defined as the difficulty of occurrence in the simulation (Supplementary Figure 1).

This method detected SNP markers as regions that may have been selected by setting the threshold to and detecting significant changes in the frequency of founder haplotypes when the value exceeds the threshold. In this study, because the focused founder haplotypes were tested using the percentage of all founder haplotypes, the horizontal axis in Supplementary Figure 1 corresponds to *k* of the multinomial distribution and the vertical axis to. Additionally, we tested the bias in each generation because we considered that founder haplotypes that increased or decreased in the early generation might not have been detected by testing the whole population. The pedigree drawing software Helium was used to separate generations (Fig. 3) ^15^. The generations were then separated into rows. Generation 1 consisted of the seven founder cultivars. Because this study compared the simulation results with actual founder haplotype frequencies, it is likely that the simulation results will be biased if the number of individuals being compared is small. Herein, the combination of columns fifth and sixth is defined as Generation 5 because there are few rows corresponding to Generation 6.

**Fig. 3.**
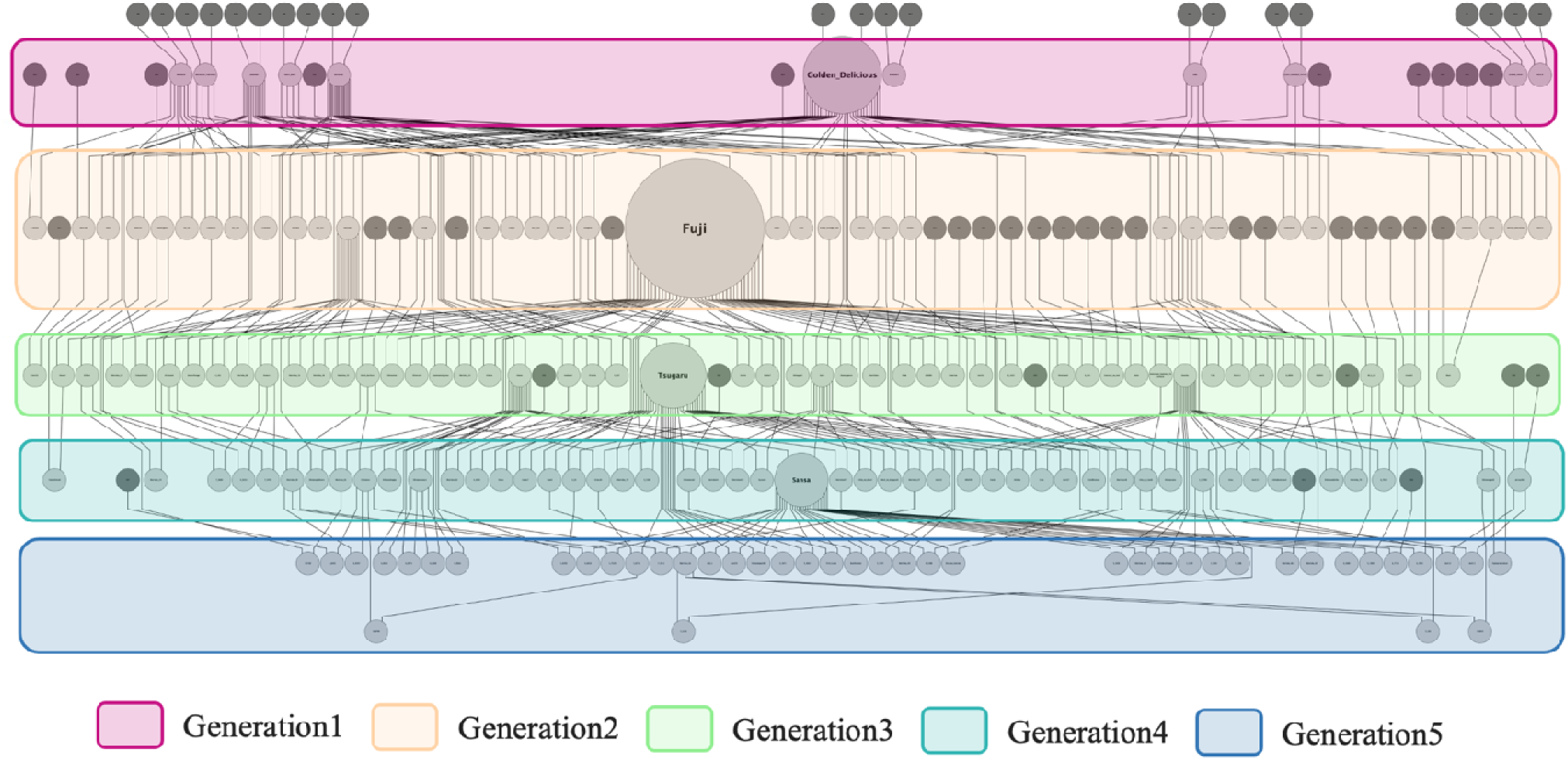
Generational partitioning of apple varieties based on the pedigree chart drawn using Helium. This diagram shows the generations and contributions of 185 Japanese varieties. Each node represents a variety, and the dark gray nodes indicate uninformed parent individuals. Each row shows the generation partitioned using the pedigree visualization tool Helium. The sizes of the nodes indicate their contribution as parents to subsequent generations. The large nodes representing ‘Fuji’ reflect its significant role in breeding, as ‘Fuji’ was used extensively for mating and has the la gest contribution to subsequent generations.

### Comparison of SNP locus, in which the frequency bias of founder haplotypes was detected, with previously reported GWAS peaks

SNP markers that significantly changed the frequency of founder haplotypes using the previous method were compared with previously reported GWAS peaks for fruit traits^14^ to determine whether the regions were selected based on fruit traits. Specifically, when the SNP marker with a significant change in founder haplotype frequency matched the SNP marker detected in the GWAS, we tracked the change in the founder haplotype frequency of the marker and the phenotypic transition by generation. Furthermore, we estimated the effects of each founder haplotype on the marker using BayesB with MCMC ^36, 37^. Marker regression models, such as BayesB, assume that genotypic values *u* are determined by a linear sum of marker effects, as follows:

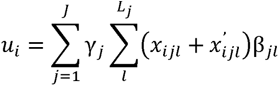

where *J* represents the total number of markers, and *L_j_* represents the number of founder haplotypes at the *j*-th marker. The variable *x_ijl_* (x*_ijl_*’) denotes the maternal (paternal) haplotype of marker *j* for variety *i* and equals to 1 if the maternal (paternal) haplotype is the *l*-th haplotype (1 = 1,2,.., *L_j_*) and 0 otherwise. The parameter *γ_j_* indicates the posterior probability of the *j*-th marker having a quantitative trait locus, while β*_jl_* represents the genetic effect associated with the *l*-th founder haplotype at marker *j*. The effect β*_jl_* is assumed to follow a Gaussian distribution 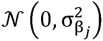, as described by Minamikawa et al.^7^. This study identified the combinations of SNP markers and founder haplotypes that exhibited significant increases or decreases in frequency. The markers were compared with GWAS peaks for fruit traits, and the set of markers that may have undergone selection based on fruit traits was narrowed. We then tracked the frequency of founder haplotype in each generation that was divided by Helium and combined with the founder haplotype effect β*_jl_*, investigated that the markers and founder haplotypes may be strongly associated with selection on fruit traits.

### Inferring reasons for changing of founder haplotype frequency via gene set enrichment analysis

The methods described above detected SNP markers showing significant changes in the frequencies of specific founder haplotypes. These markers were compared with the GWAS results for fruit traits to identify the regions potentially selected for fruit traits. However, markers that are unrelated to fruit traits may not have been detected. It is possible that these SNP markers were selected based on unmeasured traits. Therefore, we investigated SNP markers associated with these traits. First, the gene regions assigned to the ‘Golden Delicious’ doubled-haploid tree (GDDH13) reference and their annotations were downloaded from Phytozome database^38^. Using the 669 SNP markers detected using the above method, gene regions containing these markers were extracted. The extracted 271 genes were then subjected to gene set enrichment analysis using the R package “ClusterProfiler”^39^. The GO terms of these genes were analyzed in three categories: biological processes, MF, and CC.

## Acknowledgments

We thank all members of the Laboratory of Biometry and Bioinformatics of The University of Tokyo for their advice regarding this study and all members of the NARO Institute of Fruit Tree Science for maintaining the apple trees. This research 541 is supported by a grant from MAFF commissioned project study on “Smart breeding 542 technologies to Accelerate the development of new varieties toward achieving “Strategy for 543 Sustainable Food Systems, MIDORI”” and JST SPRING grant number JPMJSP2108.

## Data availability

Data supporting the findings of this study are available from the corresponding author, H.I., upon reasonable request.

## Author Contributions

The study was conceptualized by H.M., M.F.M., K.H., and H.I.. The experimental design was implemented by M.K., S.M., Y.K., and T.Y., who extracted DNA, and performed SNP genotyping. S.M. performed the phenotyping. M.F.M., M.K., and H.I. traced the founder haplotypes. M.F.M. and K.N. performed the GWAS. H.M., M.F.M., and K.H. performed gene drop simulations. H.M. and M. F. M. checked the annotation and performed GO enrichment analysis. M.F.M, K.H., and H.I. provided technical help for the statistical analysis. H.M., M.F.M., and H.I. drafted the manuscript. All the authors have read and approved the manuscript.

---

**Supplementary Figure 1.**
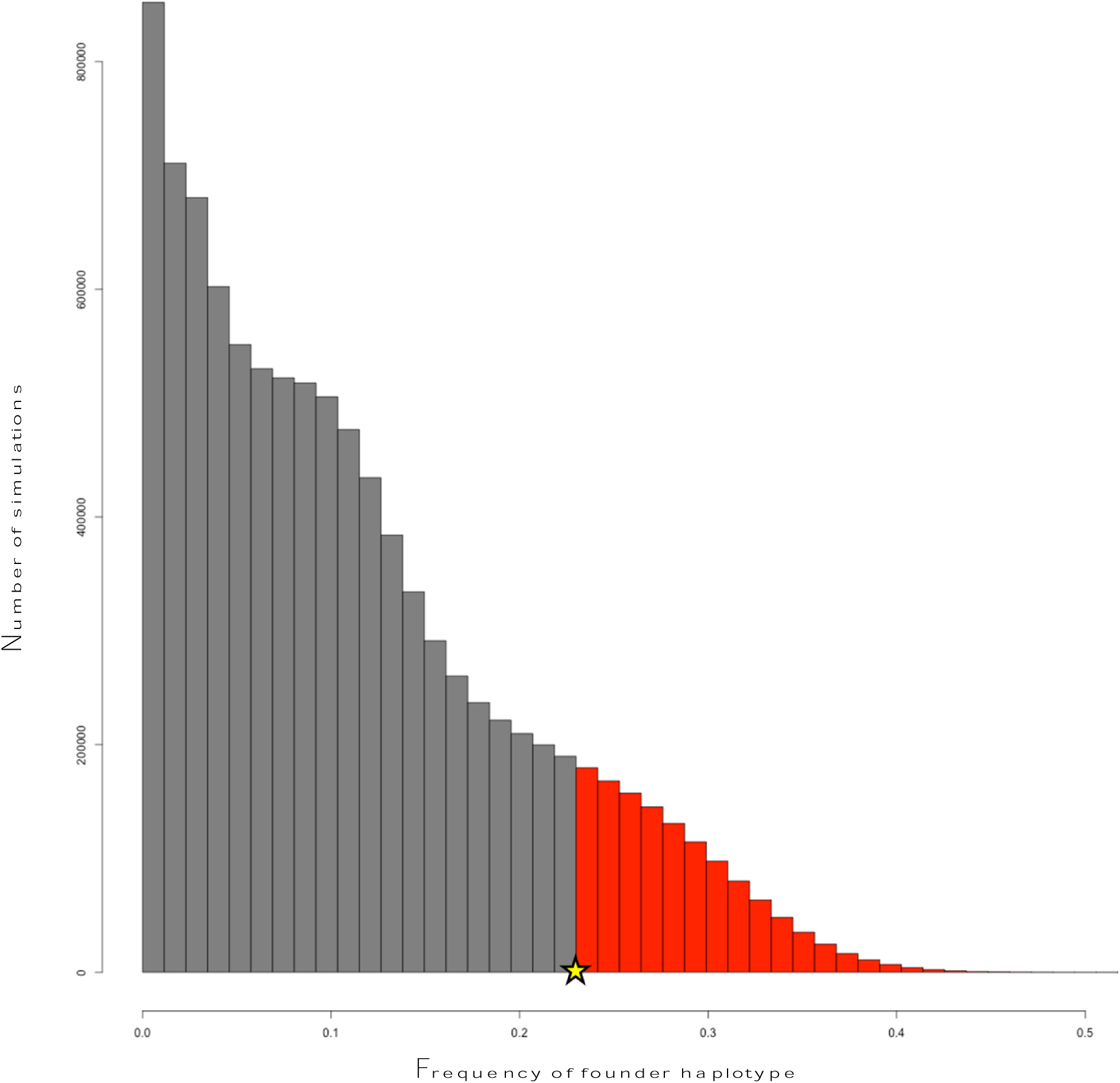
Test of Founder haplotype frequency using the null distribution. This figure shows the null distribution (grey) used to test the founder haplotype frequency. The stars indicate the position of the observed frequency of the founder haplotype at an SNP locus, and the red area shows the probability (= *p*-value) of an event being rarer than the observed frequency of the founder haplotype in the simulation.

**Supplementary Figure 2.**
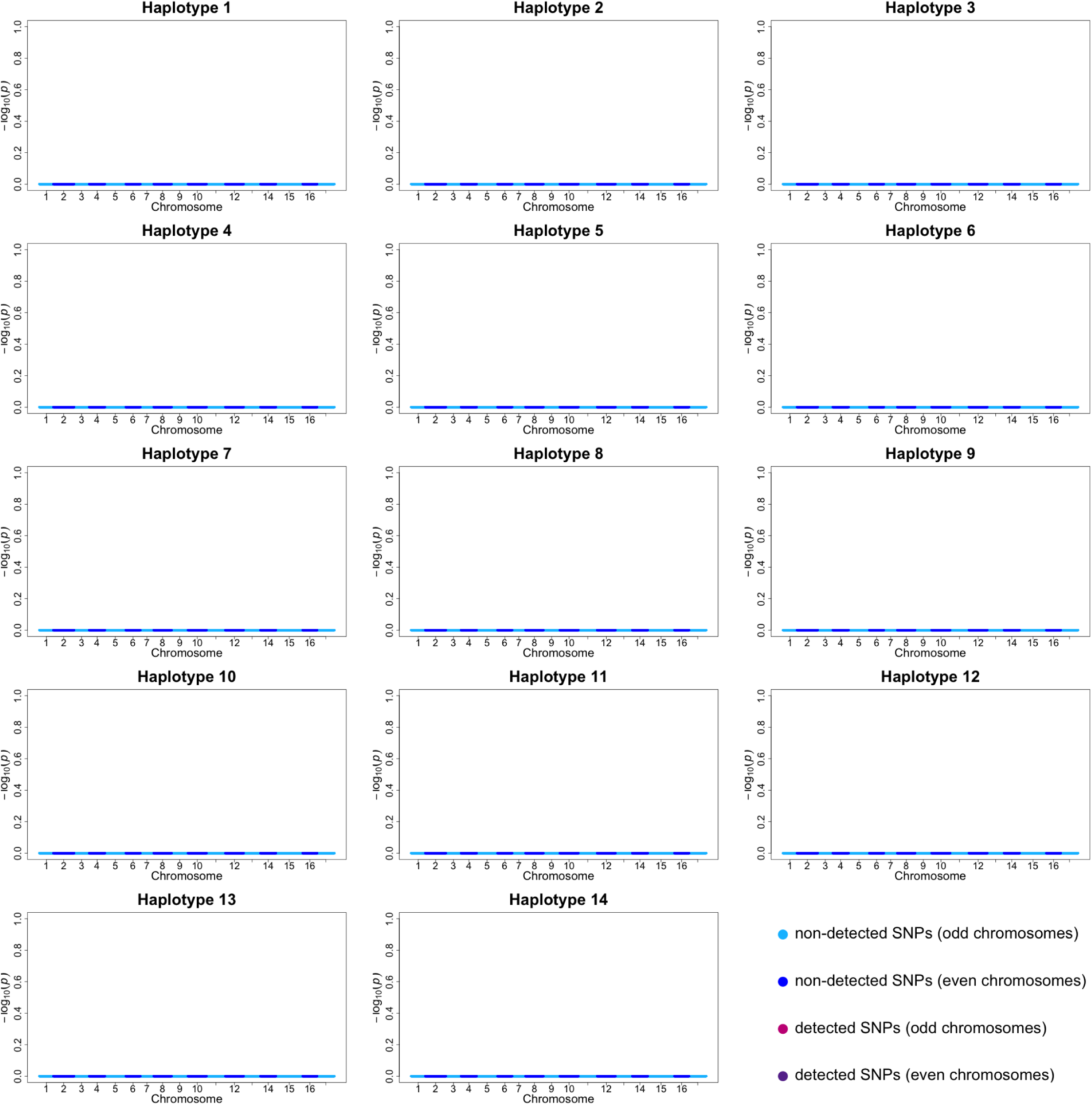
Detection of bias in the frequency of the 14 founder haplotypes in the gene drop simulation in the first generation of divided generations. No gene drop simulation was performed in the first generation because of the origin of founder haplotype.

**Supplementary Figure 3.**
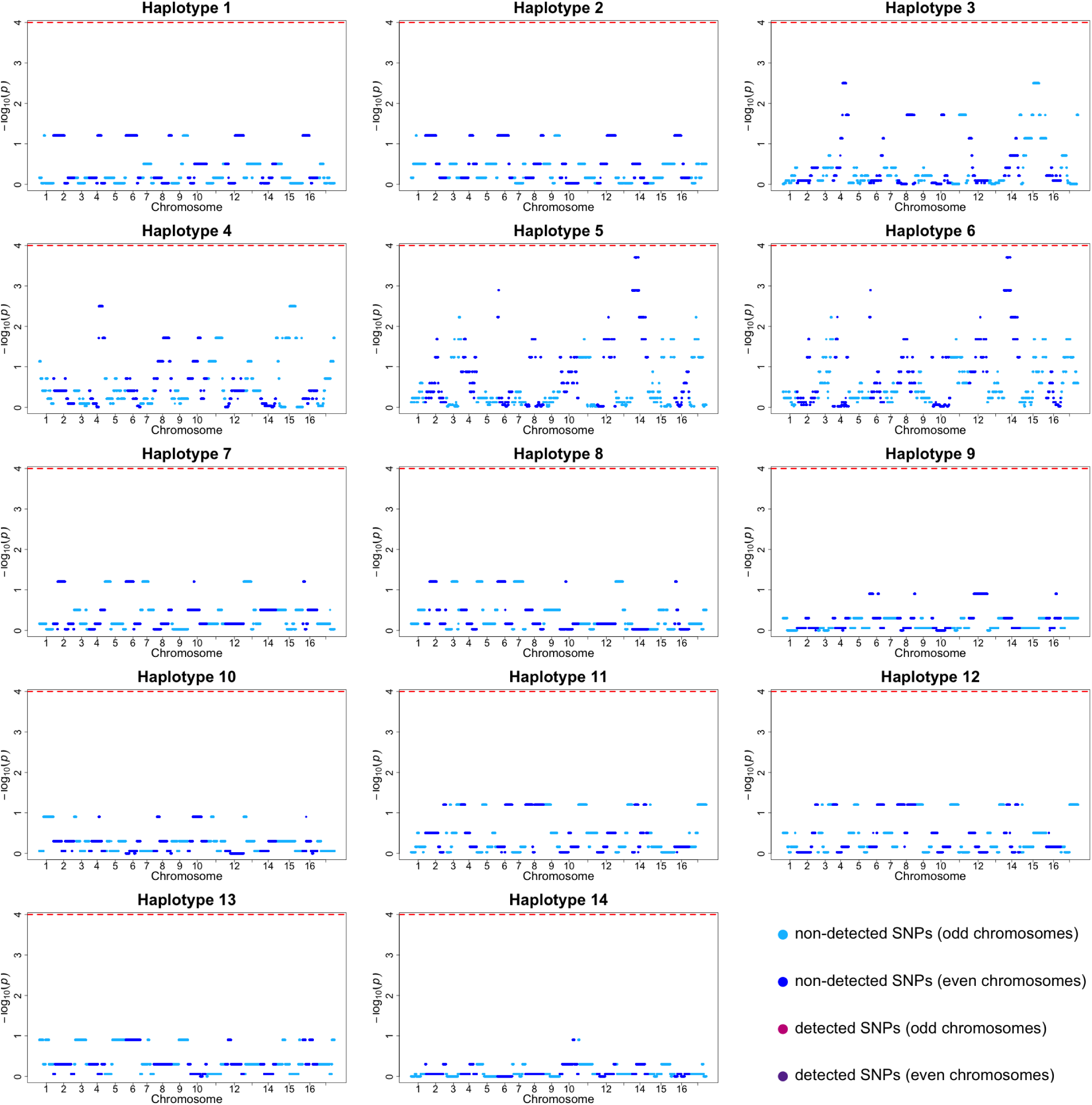
Detection of bias in the frequency of the 14 founder haplotypes in the gene drop simulation in the second generation of divided generations. Since the gene drop simulation assumes a half inheritance of each founder haplotype combination from both parents, there were no SNP markers with significantly altered founder haplotype frequencies until the second generation.

**Supplementary Figure 4.**
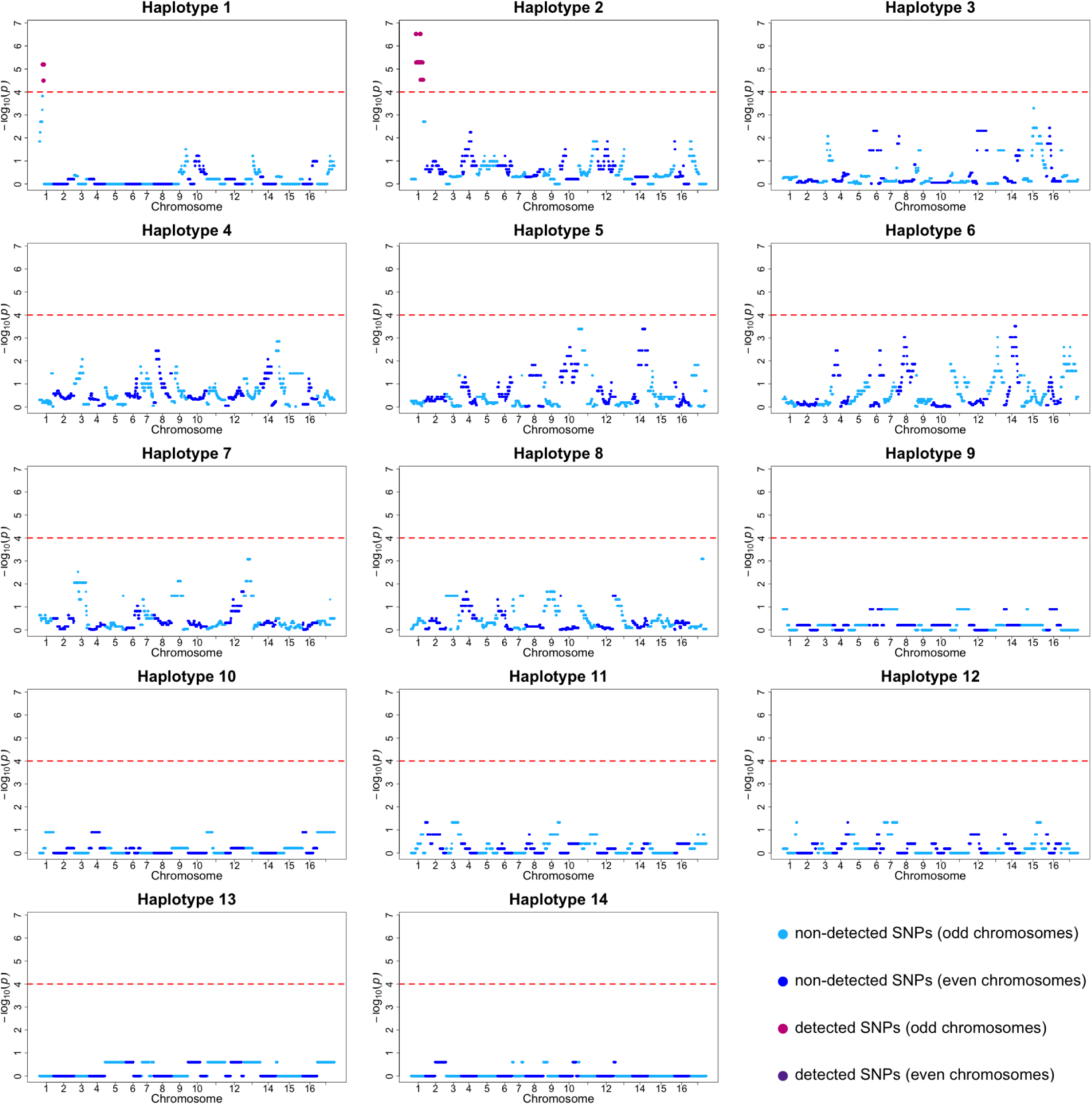
Detection of bias in the frequency of the 14 founder haplotypes in the gene drop simulation in the third generation of divided generations. SNP markers with significantly altered founder haplotype frequencies were detected in founder haplotypes 1 and 2. Both of these SNPs were located on chromosome 1.

**Supplementary Figure 5.**
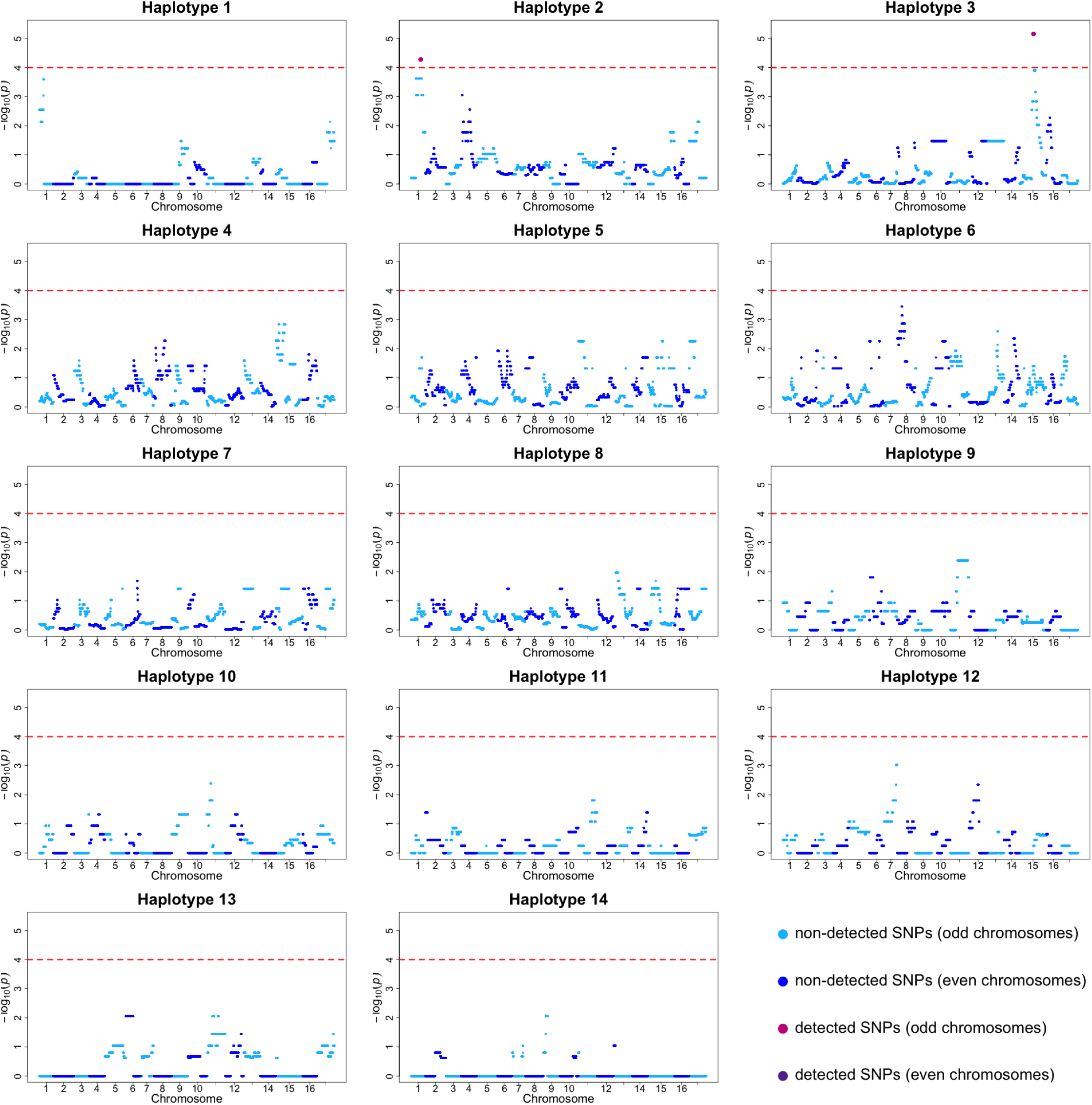
Detection of bias in the frequency of the 14 founder haplotypes in the gene drop simulation in the fourth generation of divided generations. As in the third generation, there was a SNP marker on chromosome 1 of the founder haplotype 2 that significantly changed the frequency of the founder haplotype. Moreover, the founder haplotype 3 also had significantly altered the frequency of founder haplotype associated with SNP markers on chromosome 15.

**Supplementary Figure 6.**
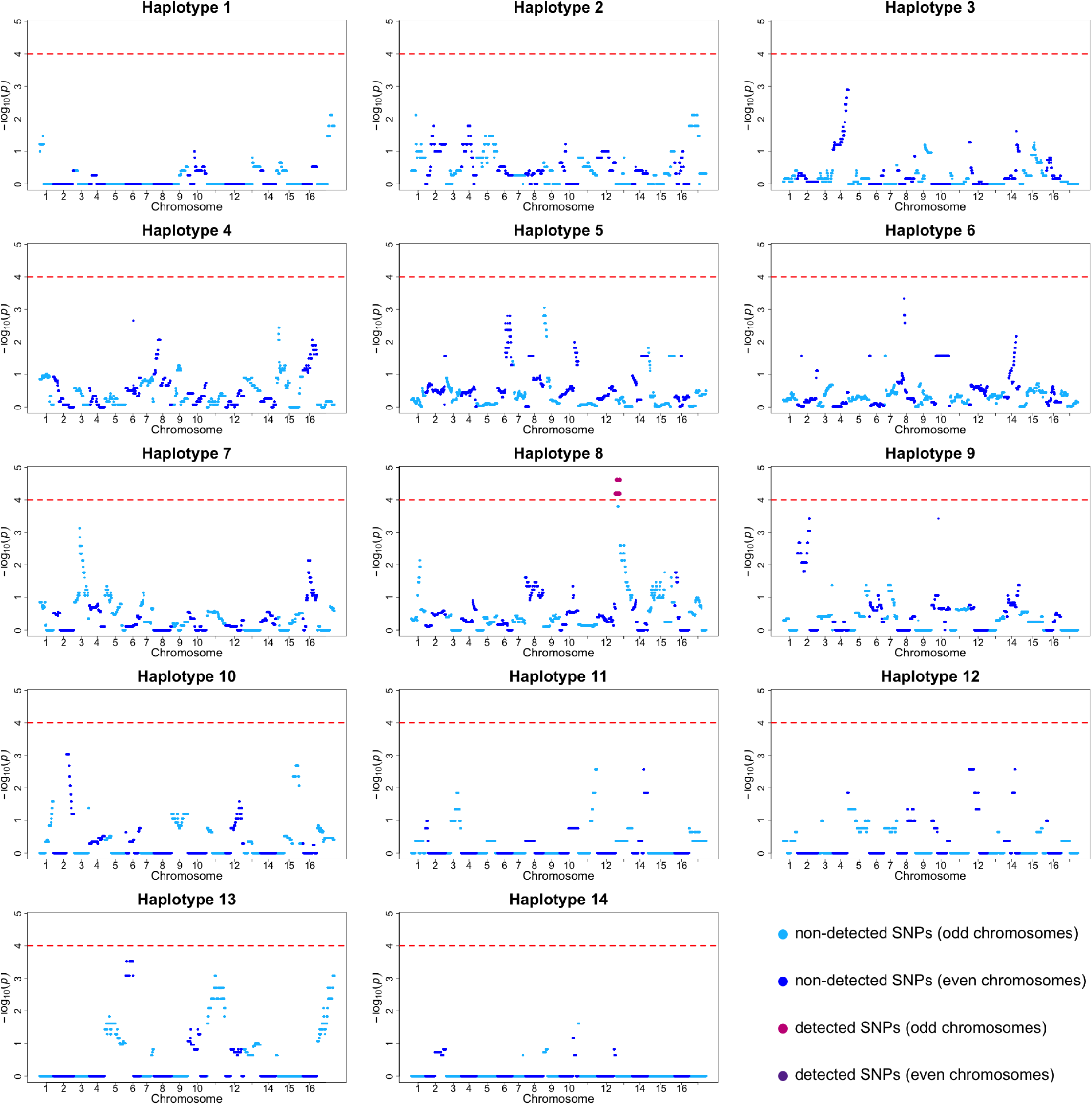
Detection of bias in the frequency of the 14 founder haplotypes in the gene drop simulation in the fifth generation of divided generations. In this generation, SNP markers significantly altering the frequency of founder haplotypes were present on chromosome 13 of founder haplotype 8.

**Supplementary Figure 7.**
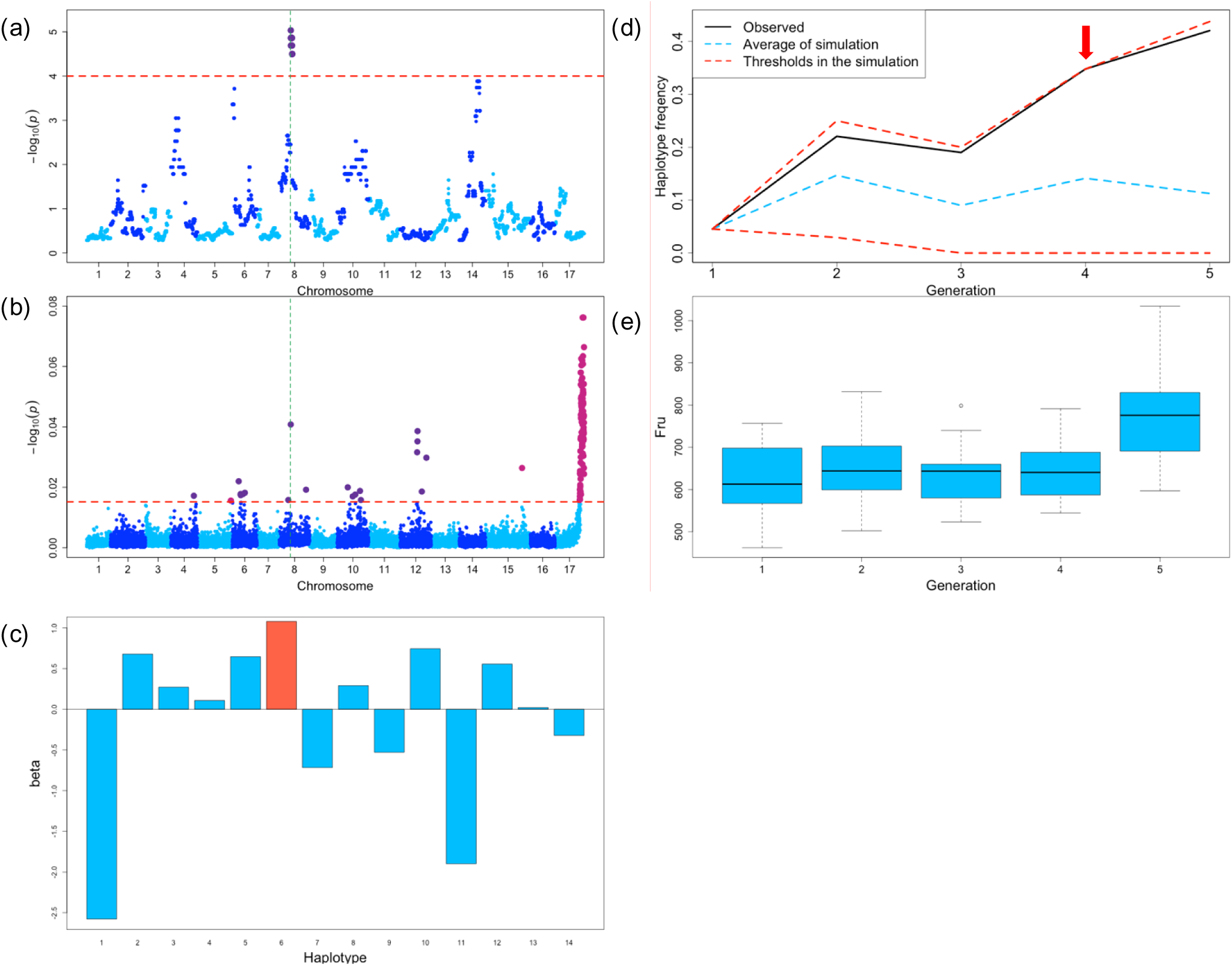
Relationship between the bias in the frequency of founder haplotype 6 from ’Golden Delicious’ and fructose content of apple fruits. (a) Detection of bias in the frequency of founder haplotype 6 from ’Golden Delicious’. (b) GWAS results for fructose content. Green dashed line in (a) and (b) indicate SNP locus (SNP_FB_1117728; block5) detected in both (a) and (b). (c) Effects (*β*) of each of the 14 founder haplotypes locating at this marker locus on fructose content. Founder haplotype 6 (red) has the largest positive effect on fructose content. (d) Changes in frequency of founder haplotype 6 against the generation. The upper and lower red dashed lines indicate the frequencies in the point of upper or lower −log_10_ (*p*) = 4, respectively, in the null distribution obtained by the simulation. The blue dashed line represents the average frequency in the simulation. (e) Box plot of fructose content in each generation. Fructose content tends to increase over generations. This marker showed an increased frequency of founder haplotype 6 with a positive effect on fructose content. There was also an increasing trend in fructose content per generation. As a result, marker effect and founder haplotype frequency matched the transition of phenotypic values.

**Supplementary Figure 8.**
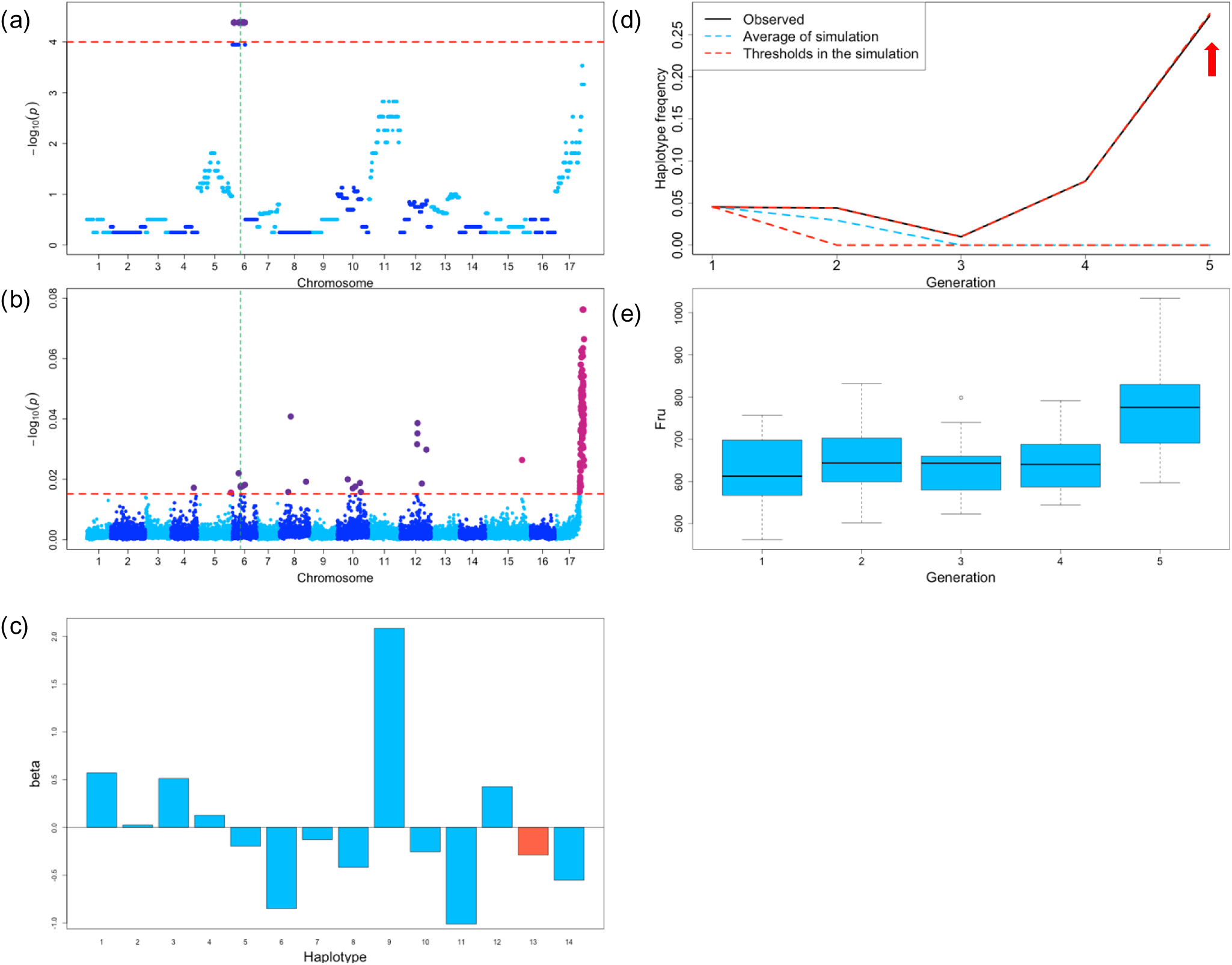
Relationship between the bias in the frequency of founder haplotype 13 from ‘Cox’s Orange Pippin’ and fructose content of apple fruits. (a) Detection of bias in the frequency of founder haplotype 13 from ‘Cox’s Orange Pippin’. (b) GWAS results for fructose content. Green dashed line in (a) and (b) indicate SNP locus (RosBREEDSNP_SNP_AG_10716913_Lg6_MDP0000302895_MAF10_MDP0000302895_exon2; block3) detected in both (a) and (b). (c) Effects (*β*) of each of the 14 founder haplotypes locating at this marker locus on fructose content. Founder haplotype 13 (red) has a fifth negative effect on fructose content. (d) Changes in frequency of founder haplotype 13 against the generation. The upper and lower red dashed lines indicate the frequencies in the point of upper or lower −log_10_ (*p*) = 4, respectively, in the null distribution obtained by the simulation. The blue dashed line represents the average frequency in the simulation. (e) Box plot of fructose content in each generation. Fructose content tends to increase over generations. This marker showed an increased frequency of founder haplotype 13 with a negative effect on fructose content. There was also an increasing trend in fructose content per generation. As a result, marker effect and frequency did not match the transition of phenotypic values.

**Supplementary Figure 9.**
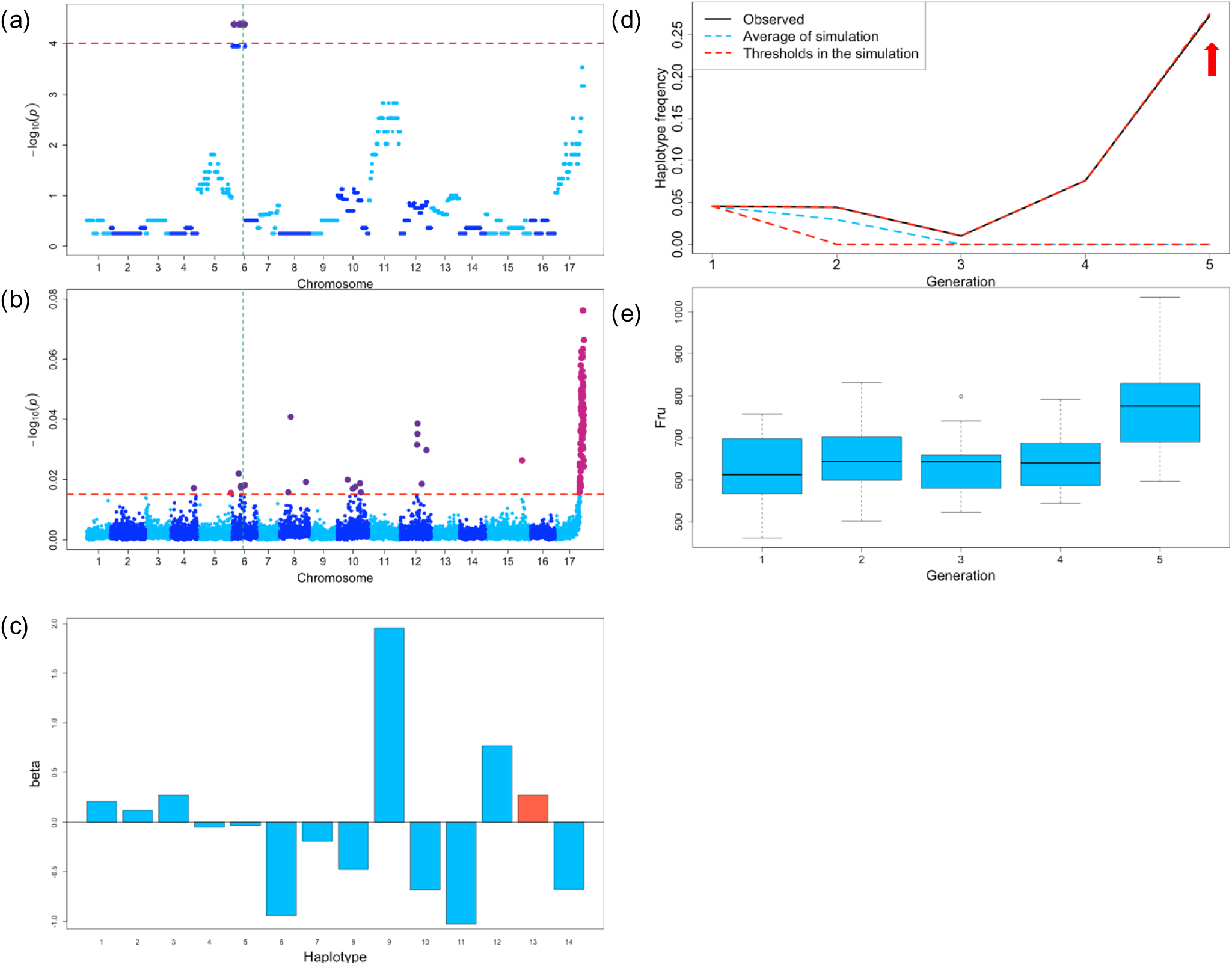
Relationship between the bias in the frequency of founder haplotype 13 from ‘Cox’s Orange Pippin’ and fructose content of apple fruits. (a) Detection of bias in the frequency of founder haplotype 13 from ‘Cox’s Orange Pippin’. (b) GWAS results for fructose content. Green dashed line in (a) and (b) indicate SNP locus (SNP_FB_0666725; block4) detected in both (a) and (b). (c) Effects (*β*) of each of the 14 founder haplotypes locating at this marker locus on fructose content. Founder haplotype 13 (red) has a third positive effect on fructose content. (d) Changes in frequency of founder haplotype 13 against the generation. The upper and lower red dashed lines indicate the frequencies in the point of upper or lower −log_10_ (*p*) = 4, respectively, in the null distribution obtained by the simulation. The blue dashed line represents the average frequency in the simulation. (e) Box plot of fructose content in each generation. Fructose content tends to increase over generations. This marker showed an increased frequency of founder haplotype 13 with a positive effect on fructose content. There was also an increasing trend in fructose content per generation. As a result, marker effect and frequency matched the transition of phenotypic values.

**Supplementary Figure 10.**
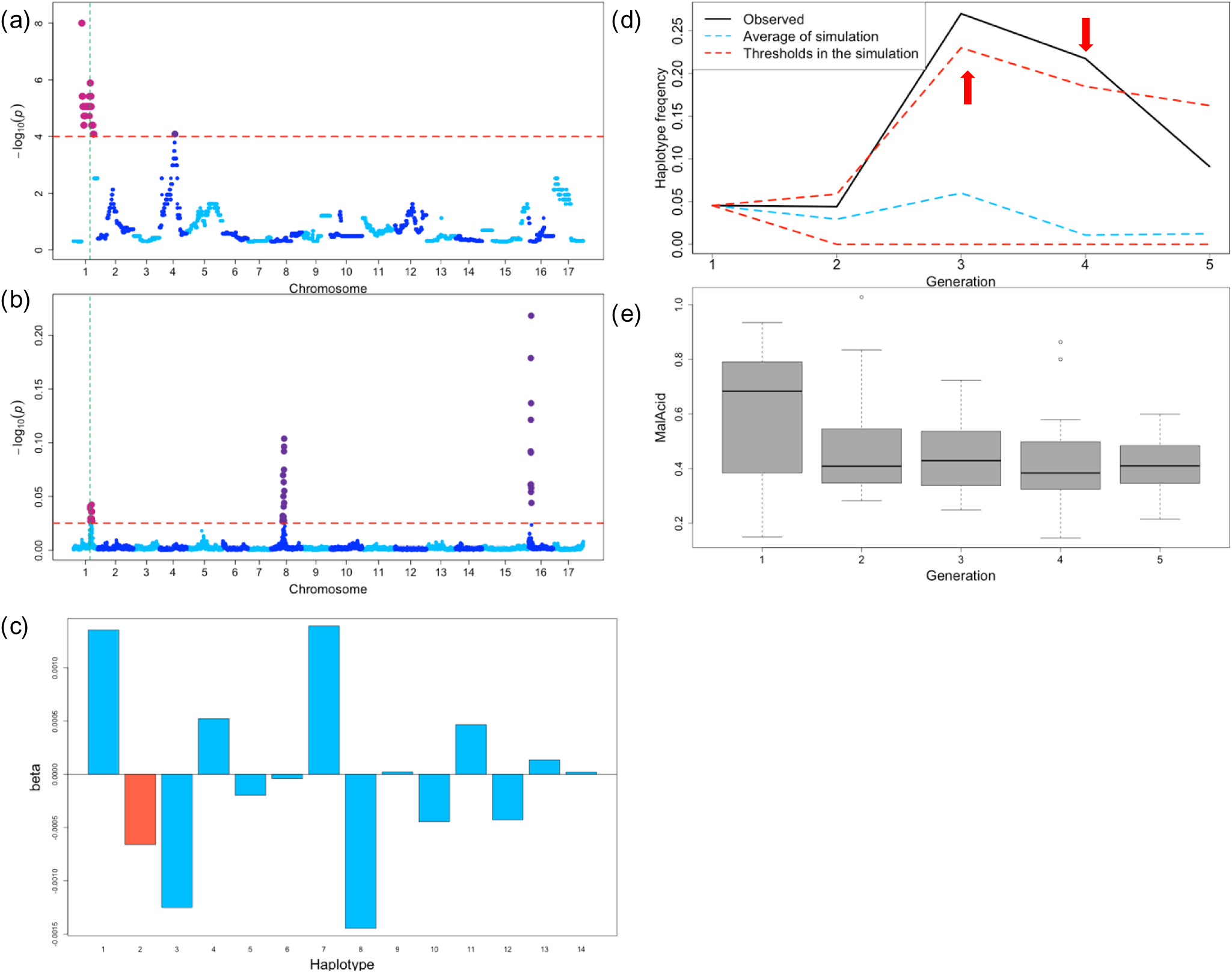
Relationship between the bias in the frequency of founder haplotype 2 from ‘Ralls Janet’ and malic acid content of apple fruits. (a) Detection of bias in the frequency of founder haplotype 2 from ‘Ralls Janet’. (b) GWAS results for malic acid content. Green dashed line in (a) and (b) indicate SNP locus (SNP_FB_0436910; block1) detected in both (a) and (b). (c) Effects (*β*) of each of the 14 founder haplotypes locating at this marker locus on malic acid content. Founder haplotype 2 (red) has a third negative effect on malic acid content. (d) Changes in frequency of founder haplotype 2 against the generation. The upper and lower red dashed lines indicate the frequencies in the point of upper or lower −log_10_ (*p*) = 4, respectively, in the null distribution obtained by the simulation. The blue dashed line represents the average frequency in the simulation. (e) Box plot of malic acid content in each generation. Malic acid content tends to decrease over generations. This marker showed an increased frequency of founder haplotype 2 with a negative effect on malic acid content. There was also an increasing trend in malic acid content per generation. As a result, marker effect and frequency matched the transition of phenotypic values.

**Supplementary Figure 11.**
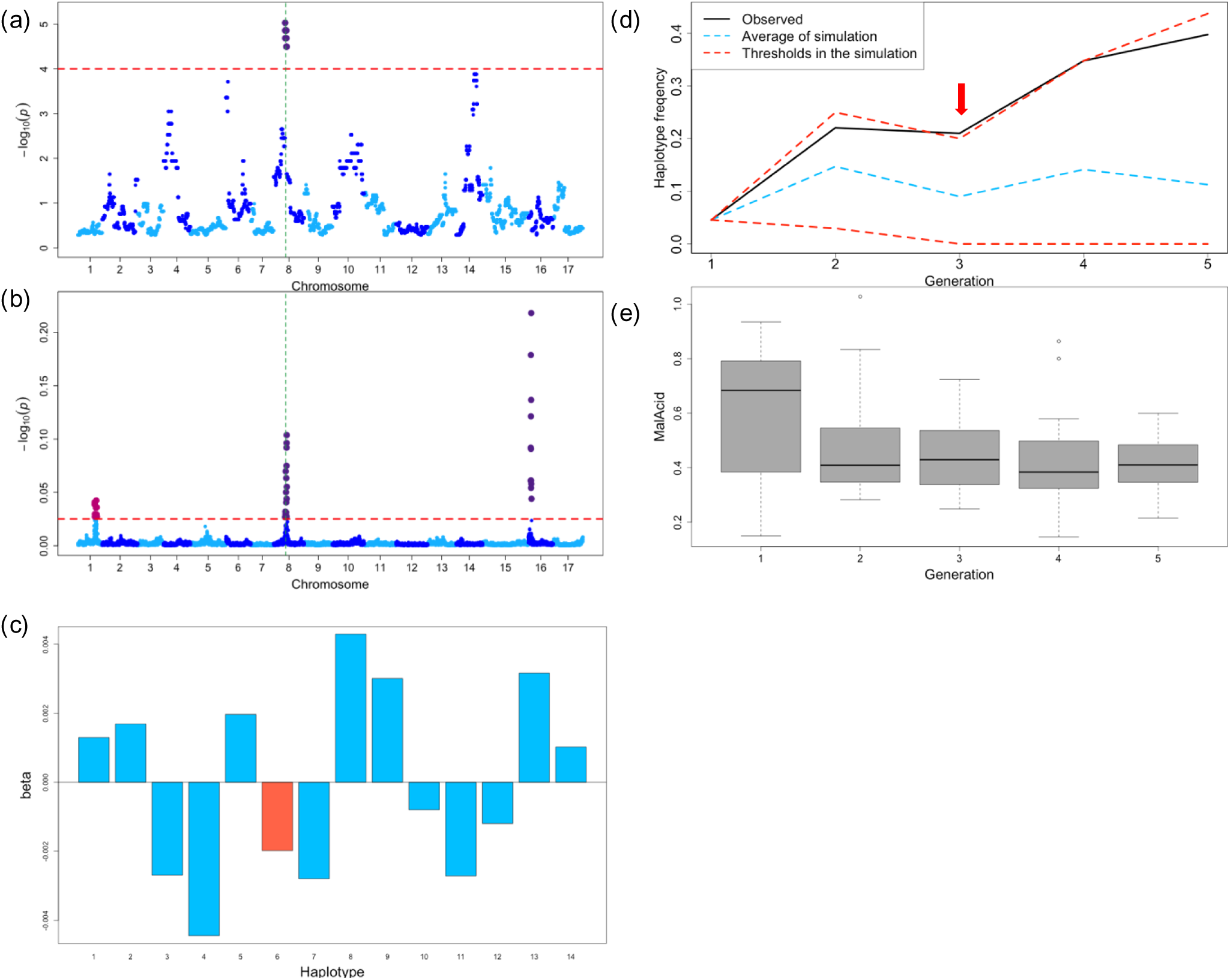
Relationship between the bias in the frequency of founder haplotype 6 from ‘Golden Delicious’ and malic acid content of apple fruits. (a) Detection of bias in the frequency of founder haplotype 6 from ‘Golden Delicious’. (b) GWAS results for malic acid content. Green dashed line in (a) and (b) indicate SNP locus (RosBREEDSNP_SNP_CT_16509379_Lg8_324685_MAF40_324685_exon1; block10) detected in both (a) and (b). (c) Effects (*β*) of each of the 14 founder haplotypes locating at this marker locus on malic acid content. Founder haplotype 6 (red) has a fifth negative effect on malic acid content. (d) Changes in frequency of founder haplotype 6 against the generation. The upper and lower red dashed lines indicate the frequencies in the point of upper or lower −log_10_ (*p*) = 4, respectively, in the null distribution obtained by the simulation. The blue dashed line represents the average frequency in the simulation. (e) Box plot of malic acid content in each generation. Malic acid content tends to decrease over generations. This marker showed an increased frequency of founder haplotype 6 with a negative effect on malic acid content. There was also a decrease trend in malic acid content per generation. As a result, marker effect and frequency matched the transition of phenotypic values.

**Supplementary Figure 12.**
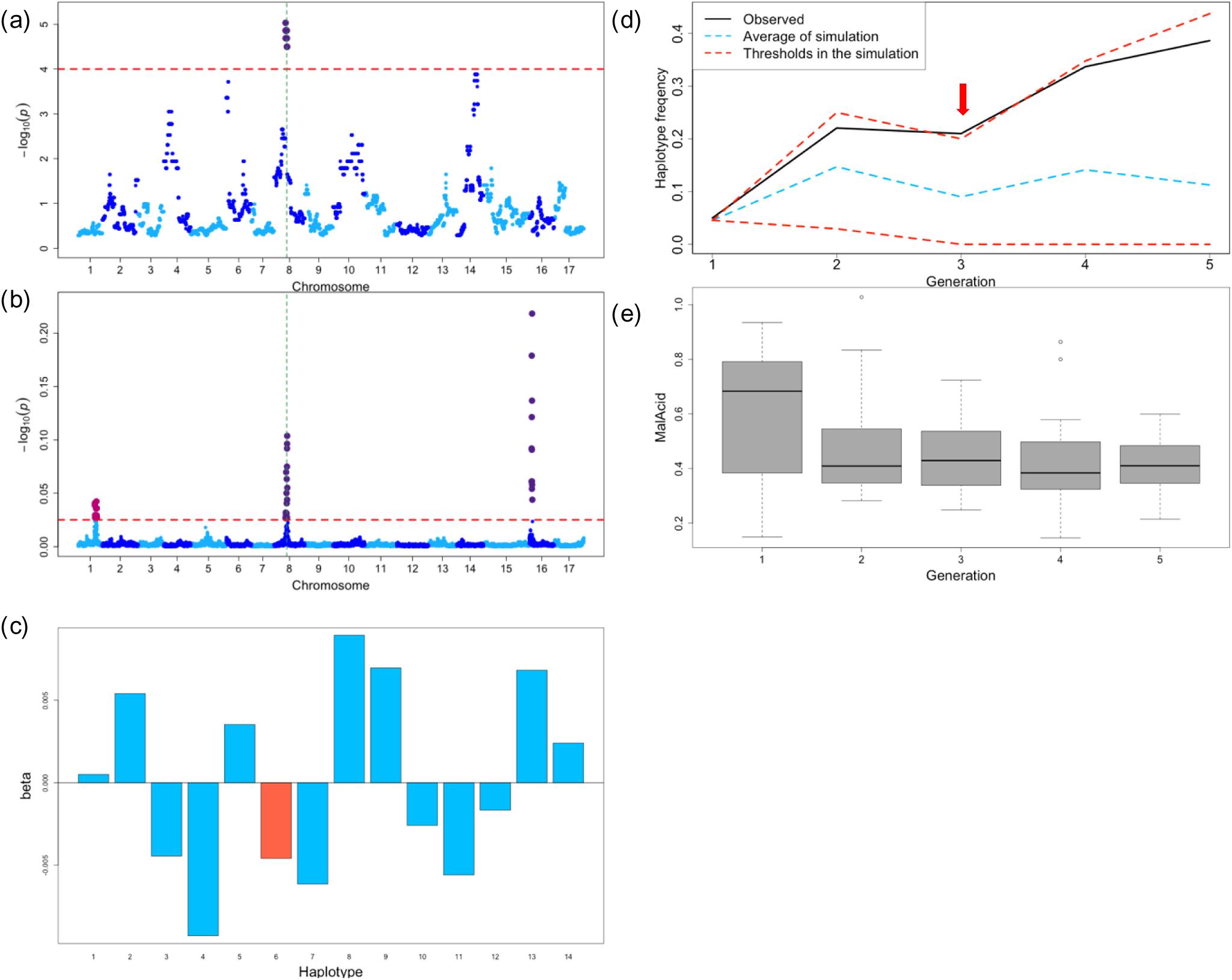
Relationship between the bias in the frequency of founder haplotype 6 from ‘Golden Delicious’ and malic acid content of apple fruits. (a) Detection of bias in the frequency of founder haplotype 6 from ‘Golden Delicious’. (b) GWAS results for malic acid content. Green dashed line in (a) and (b) indicate SNP locus (SNP_FB_0746410; block6) detected in both (a) and (b). (c) Effects (*β*) of each of the 14 founder haplotypes locating at this marker locus on malic acid content. Founder haplotype 6 (red) has a fourth negative effect on malic acid content. (d) Changes in frequency of founder haplotype 6 against the generation. The upper and lower red dashed lines indicate the frequencies in the point of upper or lower −log_10_ (*p*) = 4, respectively, in the null distribution obtained by the simulation. The blue dashed line represents the average frequency in the simulation. (e) Box plot of malic acid content in each generation. Malic acid content tends to decrease over generations. This marker showed an increased frequency of founder haplotype 6 with a negative effect on malic acid content. There was also a decrease trend in malic acid content per generation. As a result, marker effect and frequency matched the transition of phenotypic values.

**Supplementary Figure 13.**
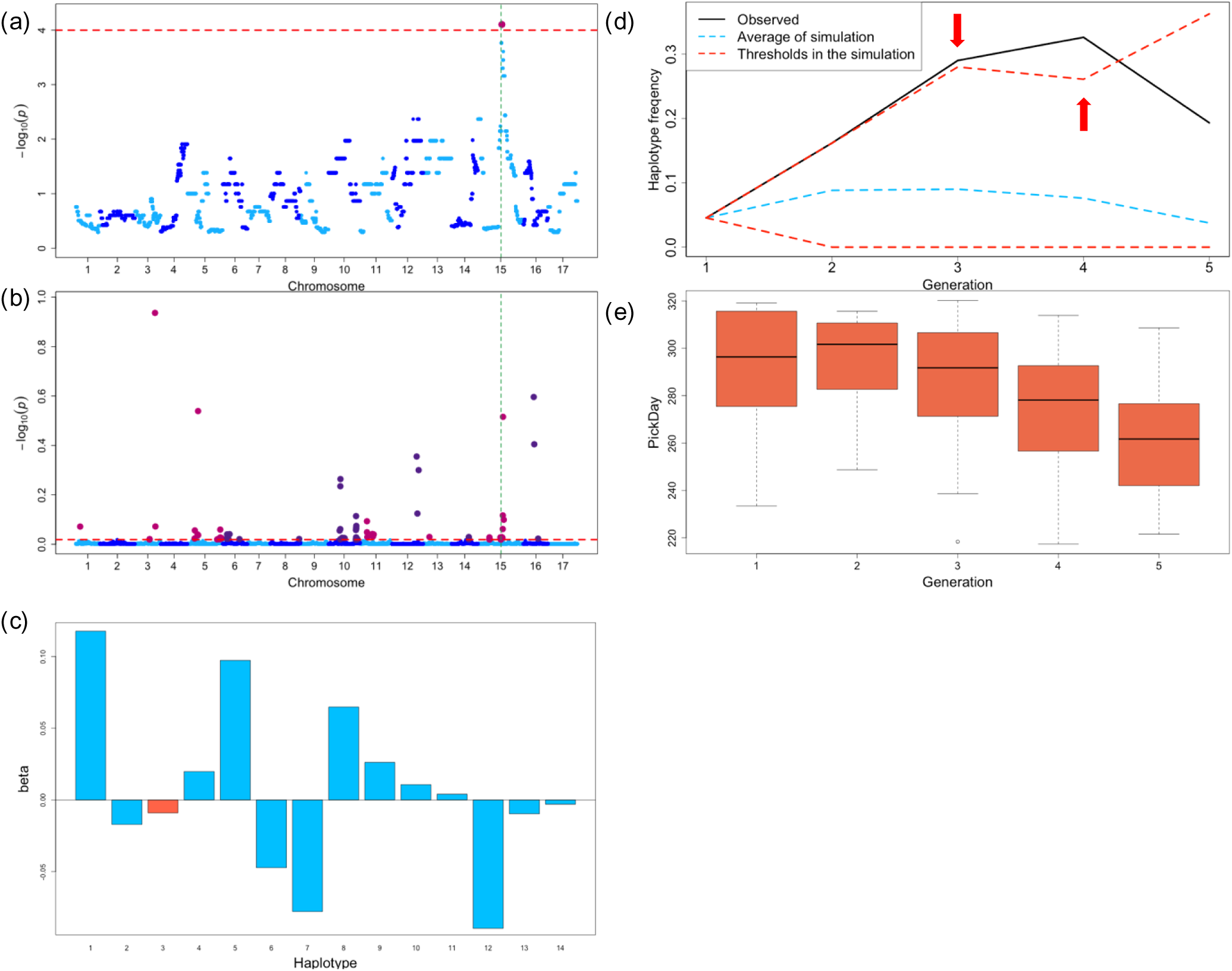
Relationship between the bias in the frequency of founder haplotype 3 from ‘Delicious’ and Harvest time (PickDay) of apple fruits. (a) Detection of bias in the frequency of founder haplotype 3 from ‘Delicious’. (b) GWAS results for PickDay. Green dashed line in (a) and (b) indicate SNP locus (SNP_FB_0303417; block11) detected in both (a) and (b). (c) Effects (*β*) of each of the 14 founder haplotypes locating at this marker locus on PickDay. Founder haplotype 3 (red) has a sixth negative effect on PickDay. (d) Changes in frequency of founder haplotype 3 against the generation. The upper and lower red dashed lines indicate the frequencies in the point of upper or lower −log_10_ (*p*) = 4, respectively, in the null distribution obtained by the simulation. The blue dashed line represents the average frequency in the simulation. (e) Box plot of PickDay in each generation. PickDay tends to decrease over generations. This marker showed an increased frequency of founder haplotype 3 with a negative effect on PickDay. There was also a decrease trend in PickDay per generation. As a result, marker effect and frequency matched the transition of phenotypic values.

**Supplementary Figure 14.**
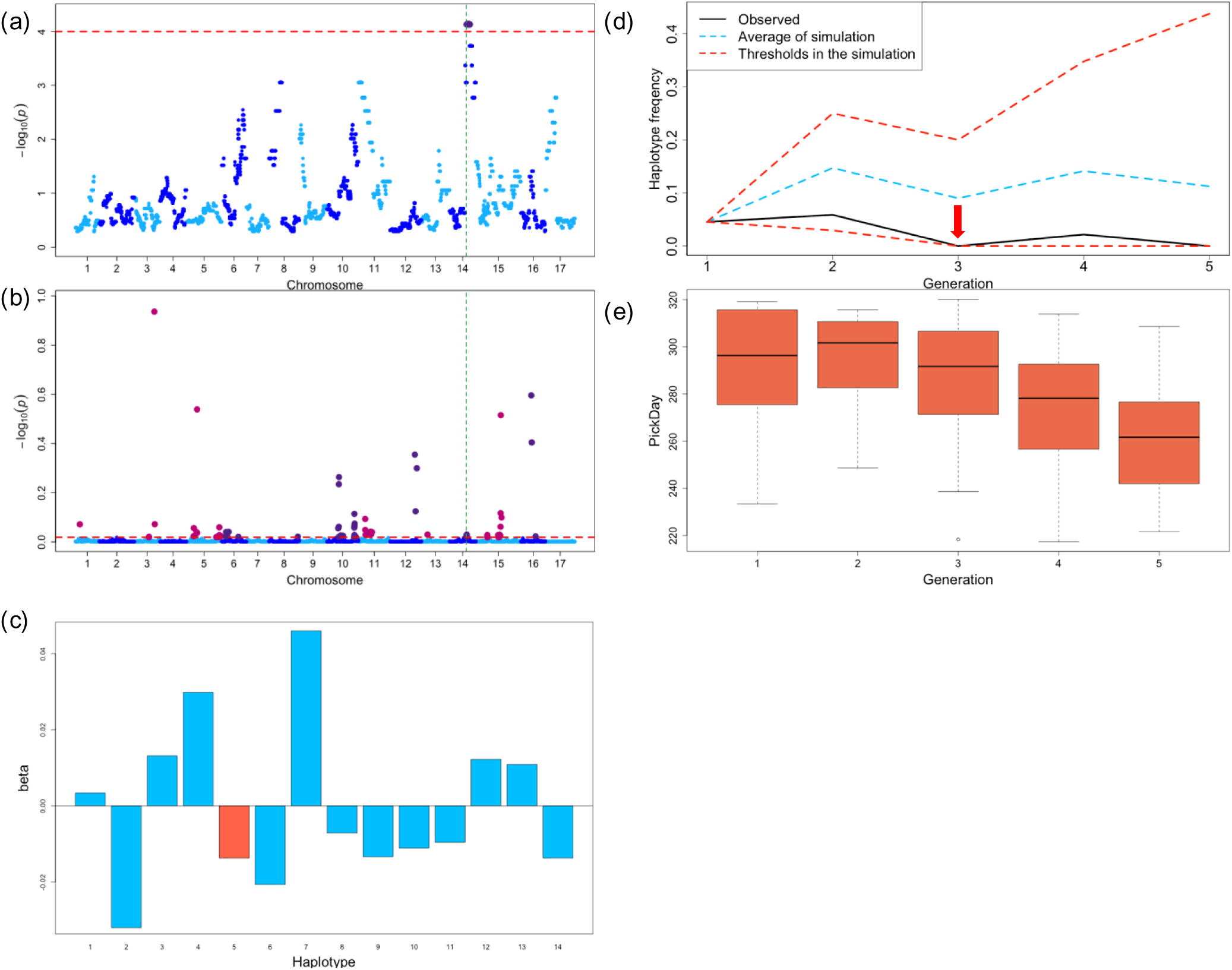
Relationship between the bias in the frequency of founder haplotype 5 from ‘Golden Delicious’ and PickDay of apple fruits. (a) Detection of bias in the frequency of founder haplotype 5 from ‘Golden Delicious’. (b) GWAS results for PickDay. Green dashed line in (a) and (b) indicate SNP locus (SNP_FB_0246031; block9) detected in both (a) and (b). (c) Effects (*β*) of each of the 14 founder haplotypes locating at this marker locus on PickDay. Founder haplotype 5 (red) has a third negative effect on PickDay. (d) Changes in frequency of founder haplotype 3 against the generation. The upper and lower red dashed lines indicate the frequencies in the point of upper or lower −log_10_ (*p*) = 4, respectively, in the null distribution obtained by the simulation. The blue dashed line represents the average frequency in the simulation. (e) Box plot of PickDay in each generation. PickDay tends to decrease over generations. This marker showed a decrease frequency of founder haplotype 5 with a negative effect on PickDay. There was also a decrease trend in PickDay per generation. As a result, marker effect and frequency did not match the transition of phenotypic values.

**Supplementary Figure 15.**
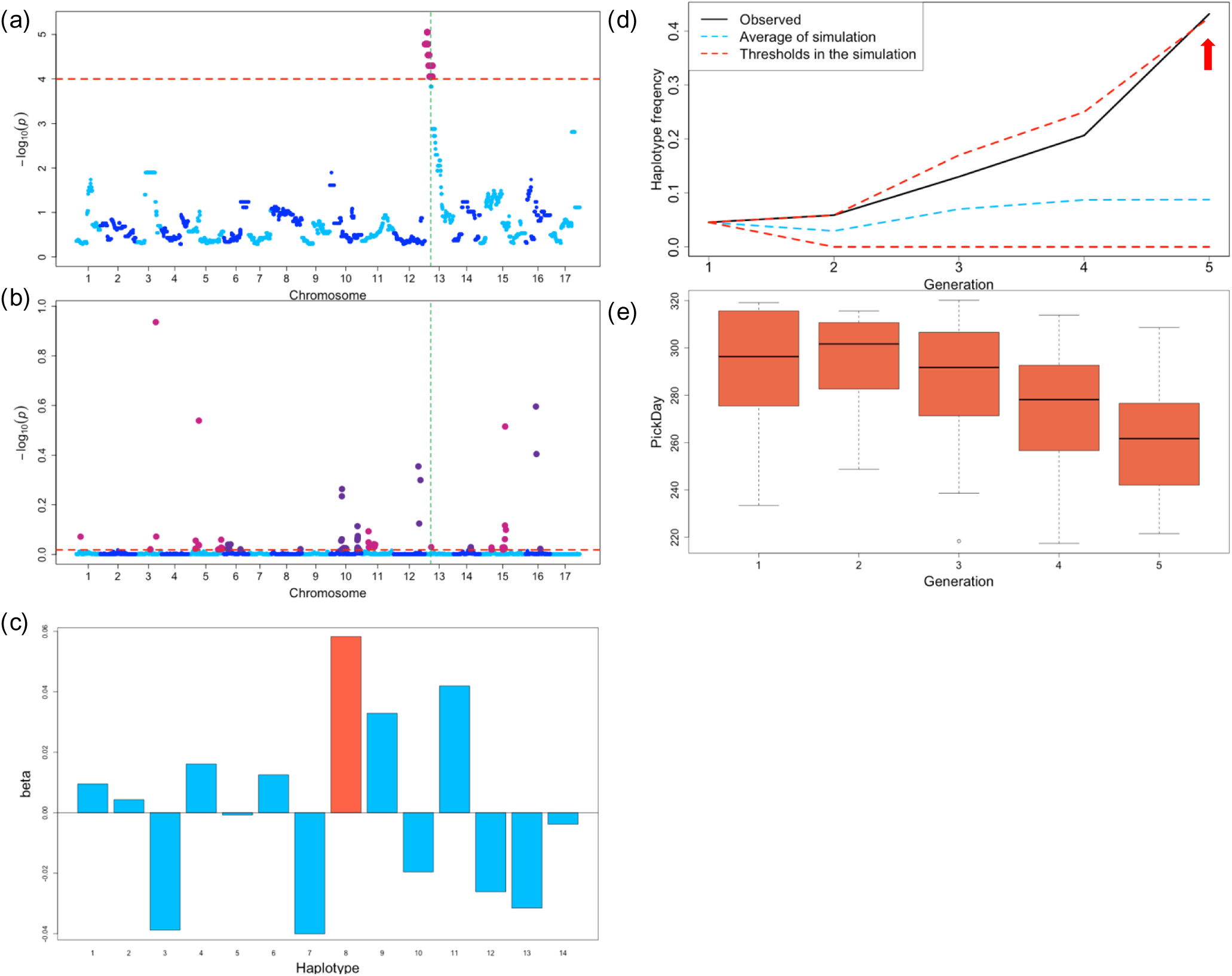
Relationship between the bias in the frequency of founder haplotype 8 from ‘Jonathan’ and PickDay of apple fruits. (a) Detection of bias in the frequency of founder haplotype 8 from ‘Jonathan’. (b) GWAS results for PickDay. Green dashed line in (a) and (b) indicate SNP locus (SNP_FB_0164463; block8) detected in both (a) and (b). (c) Effects (*β*) of each of the 14 founder haplotypes locating at this marker locus on PickDay. Founder haplotype 8 (red) has a largest positive effect on PickDay. (d) Changes in frequency of founder haplotype 8 against the generation. The upper and lower red dashed lines indicate the frequencies in the point of upper or lower −log_10_ (*p*) = 4, respectively, in the null distribution obtained by the simulation. The blue dashed line represents the average frequency in the simulation. (e) Box plot of PickDay in each generation. PickDay tends to decrease over generations. This marker showed an increase frequency of founder haplotype 8 with a positive effect on PickDay. There was also a decrease trend in PickDay per generation. As a result, marker effect and frequency did not match the transition of phenotypic values.

**Supplementary Figure 16.**
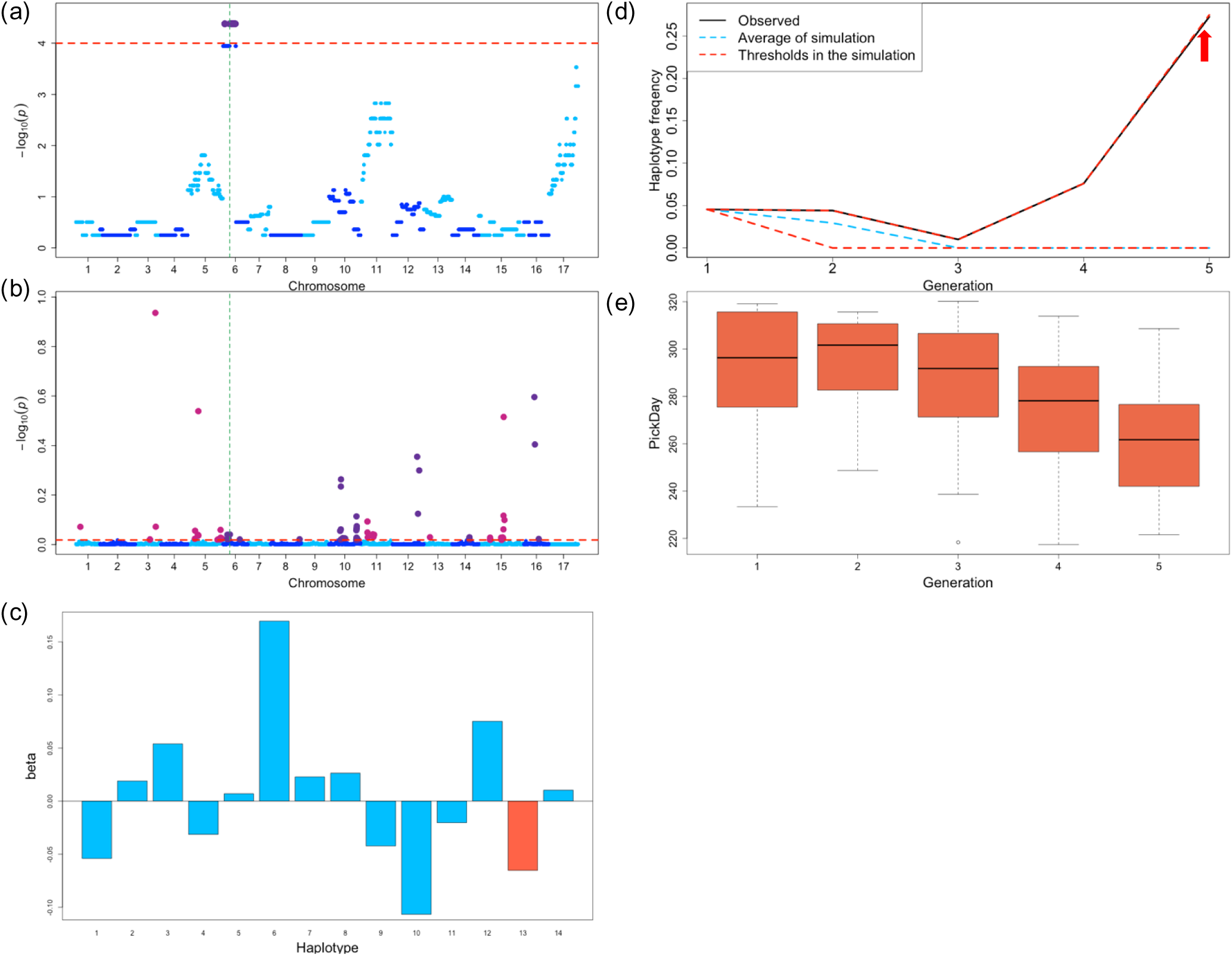
Relationship between the bias in the frequency of founder haplotype 13 from ‘Cox’s Orange Pippin’ and PickDay of apple fruits. (a) Detection of bias in the frequency of founder haplotype 13 from ‘Cox’s Orange Pippin’. (b) GWAS results for PickDay. Green dashed line in (a) and (b) indicate SNP locus (SNP_FB_0654632; block2) detected in both (a) and (b). (c) Effects (*β*) of each of the 14 founder haplotypes locating at this marker locus on PickDay. Founder haplotype 13 (red) has a second negative effect on PickDay. (d) Changes in frequency of founder haplotype 13 against the generation. The upper and lower red dashed lines indicate the frequencies in the point of upper or lower −log_10_ (*p*) = 4, respectively, in the null distribution obtained by the simulation. The blue dashed line represents the average frequency in the simulation. (e) Box plot of PickDay in each generation. PickDay tends to decrease over generations. This marker showed an increase frequency of founder haplotype 13 with a negative effect on PickDay. There was also a decrease trend in PickDay per generation. As a result, marker effect and frequency matched the transition of phenotypic values.

**Supplementary Figure 17.**
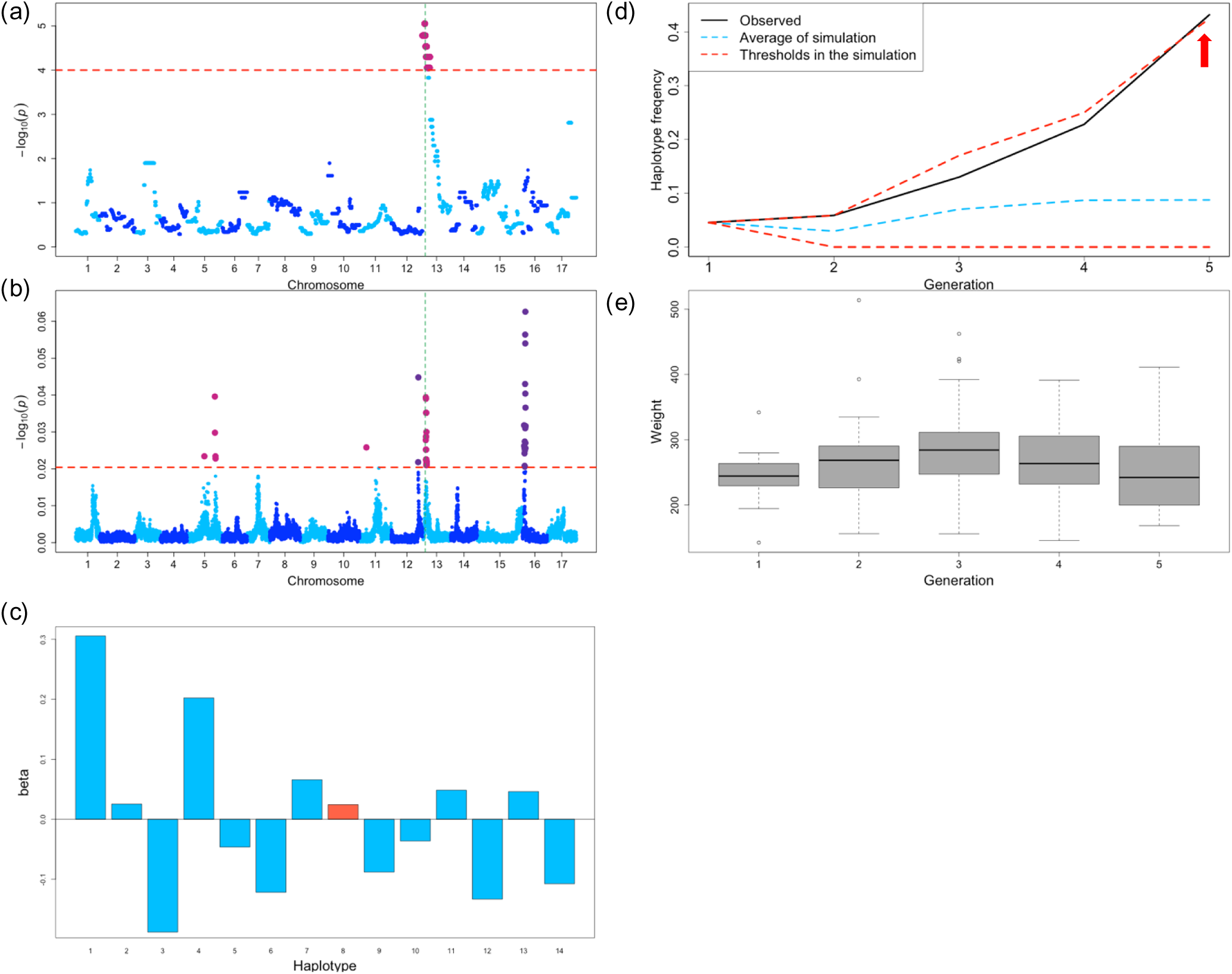
Relationship between the bias in the frequency of founder haplotype 8 from ‘Jonathan’ and fruit weight (Weight) of apple fruits. (a) Detection of bias in the frequency of founder haplotype 8 from ‘Jonathan’. (b) GWAS results for Weight. Green dashed line in (a) and (b) indicate SNP locus (SNP_FB_0160784; block7) detected in both (a) and (b). (c) Effects (*β*) of each of the 14 founder haplotypes locating at this marker locus on Weight. Founder haplotype 8 (red) has a seventh positive effect on Weight. (d) Changes in frequency of founder haplotype 8 against the generation. The upper and lower red dashed lines indicate the frequencies in the point of upper or lower −log_10_ (*p*) = 4, respectively, in the null distribution obtained by the simulation. The blue dashed line represents the average frequency in the simulation. (d) Box plot of Weight in each generation. Weight has no trend over generations. This marker showed an increase frequency of founder haplotype 8 with a positive effect on Weight. There was also a no trend in Weight per generation. As a result, marker effect and frequency did not match the transition of phenotypic values.

**Supplementary Figure 18.**
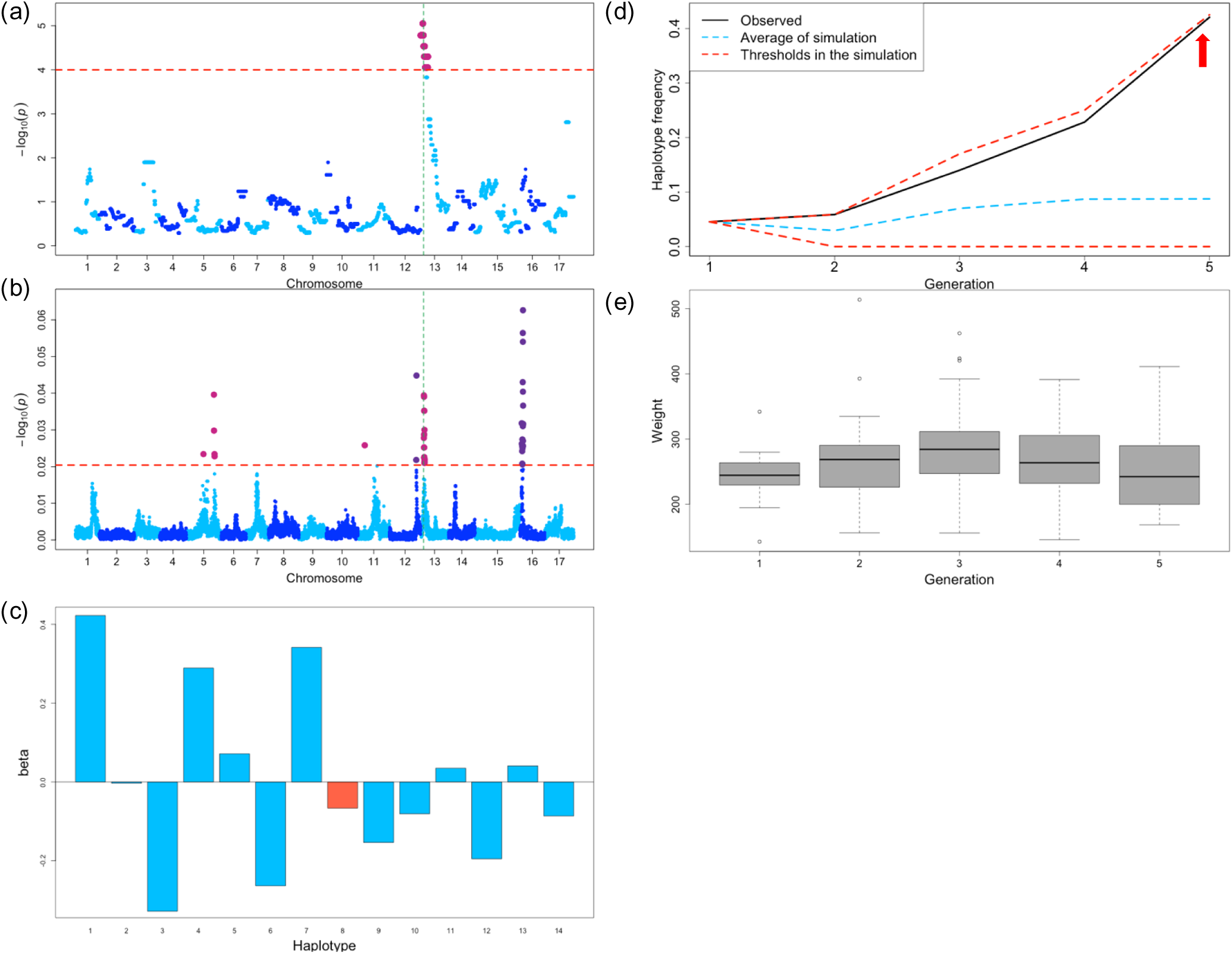
Relationship between the bias in the frequency of founder haplotype 8 from ‘Jonathan’ and Weight of apple fruits. (a) Detection of bias in the frequency of founder haplotype 8 from ‘Jonathan’. (b) GWAS results for Weight. Green dashed line in (a) and (b) indicate SNP locus (SNP_FB_1063136; block12) detected in both (a) and (b). (c) Effects (*β*) of each of the 14 founder haplotypes locating at this marker locus on Weight. Founder haplotype 8 (red) has a seventh negative effect on Weight. (d) Changes in frequency of founder haplotype 8 against the generation. The upper and lower red dashed lines indicate the frequencies in the point of upper or lower −log_10_ (*p*) = 4, respectively, in the null distribution obtained by the simulation. The blue dashed line represents the average frequency in the simulation. (e) Box plot of Weight in each generation. Weight has no trend over generations. This marker showed an increase frequency of founder haplotype 8 with a negative effect on Weight. There was also a no trend in Weight per generation. As a result, marker effect and frequency did not match the transition of phenotypic values.

**Supplementary Figure 19.**
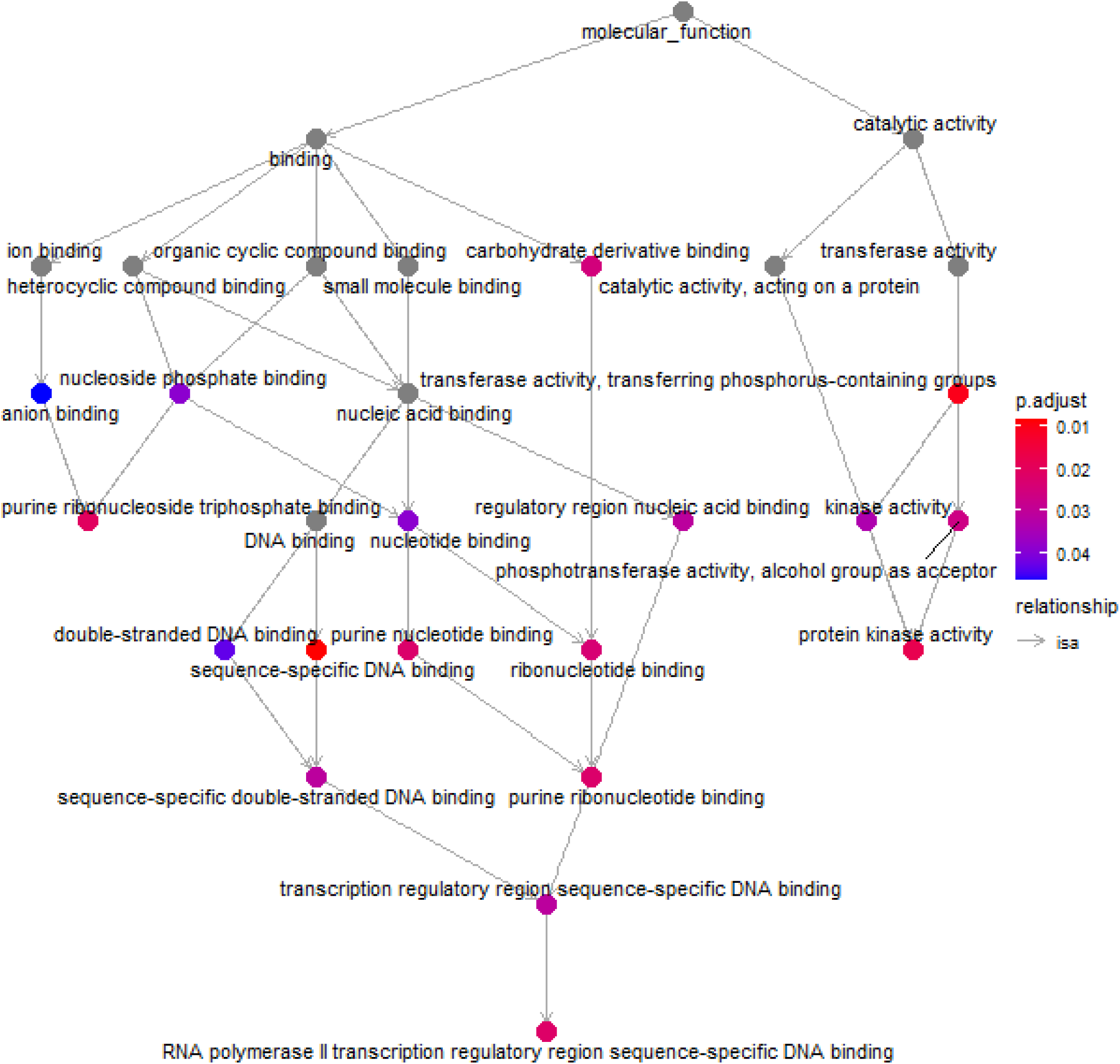
MF gene network drawing by go-plot in ClusterProfiler The result of gene set enrichment analysis in MF. Arrows indicate relationship between functions. MF had two type of function and its separated binding and catalytic activity. In catalytic activity, the transferase activity, transferring phosphorus-containing groups had lowest adjusted p-value. Additionally, the protein kinase activity is most down-stream function. Moreover, in binding function, sequence-specific DNA binding had lowest adjusted p-value. The most down-stream function of binding is RNA polymerase II transcription regulatory region sequence-specific DNA binding.

**Supplementary Figure 20.**
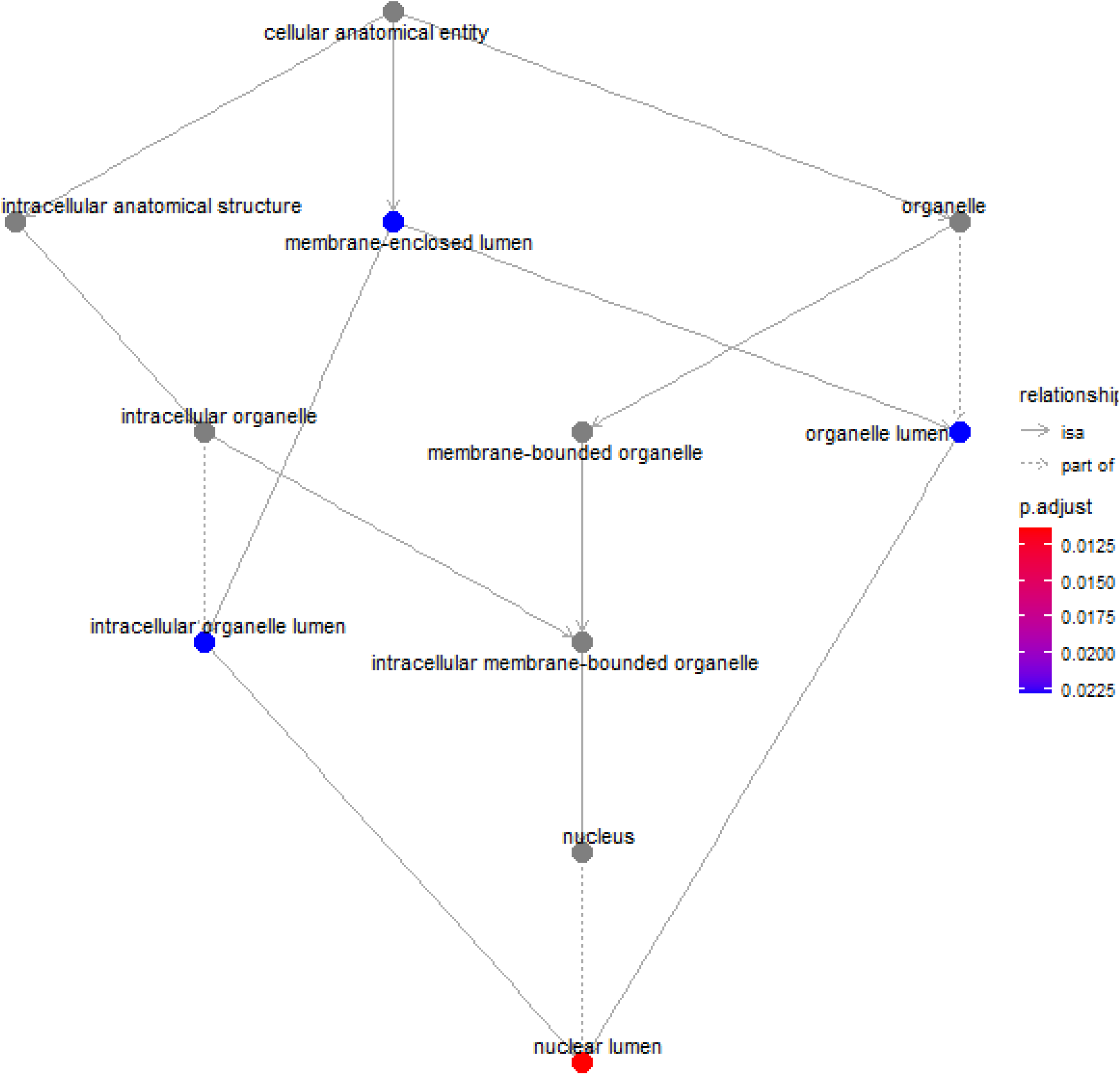
CC gene network drawing by go-plot in ClusterProfiler The result of gene set enrichment analysis in CC. The solid arrows indicate relationship between functions and the dashed arrows indicate part of the function is included. The function related to nuclear lumen is most down-stream and significant. The function CC related to the SNP markers, associated with biased founder haplotype frequency, may be related to nuclear lumen.

